# Effect of different odors on the rat urine proteome

**DOI:** 10.1101/2024.04.17.590001

**Authors:** Yuqing Liu, Haitong Wang, Youhe Gao

**Affiliations:** (Gene Engineering Drug and Biotechnology Beijing Key Laboratory, College of Life Sciences Beijing Normal University, Beijing 100875, China)

**Keywords:** Urine, Proteomics, Odor

## Abstract

Do rats have corresponding changes in their urinary proteome when smelling different odors? In this study, urine samples were collected from six rats after smelling sesame oil and essential balm for three days. And samples were collected before and on the third and fourth days. Comparing the urinary protein groups of Day0 and Day4 of the sesame oil group, 143 differential proteins were identified, and the average number of randomly generated differential proteins was 7.3, which means that about 95% of the differential proteins could not be randomly generated. in the sesame oil group, differential proteins such as low-density lipoprotein receptor-related protein 2 and fetuin B, a biomarker of COPD, which are associated with olfaction, were identified. While uteroglobulin, trichothecene factor 3, and visfatin 2 were identified in the essential balm group, which had significant changes and were related to the production of olfactory sensation. It is noteworthy that we identified odor-binding protein 2A in the essential balm group, which was present in the e-cigarette model. This study demonstrates that odor can affect rat urinary proteome, with different odors affecting it differently. This provides a new approach to explore the biological process of olfaction.

## 1. Introduction

### 1.1 Urine Biomarkers

Biomarkers are indicators that can objectively reflect normal pathological processes as well as physiological processes^[1]^, and clinically, biomarkers can predict, monitor, and diagnose multifactorial diseases at different stages^[2]^. The potential of urinary biomarkers has not been fully developed compared to more widely used blood biomarkers. However, urinary biomarkers are especially important in terms of early diagnosis of disease and prediction of status. Since homeostatic mechanisms are regulated in the blood, changes in the blood proteome caused by disease are metabolically excreted, and no significant changes can be apparent in the early stages of the disease. Whereas urine is produced by glomerular filtration of plasma and is not regulated by homeostatic mechanisms, and minor changes in the disease at an early stage can be observed in urine. According to a previous study, the changes observed in urine occur far earlier than those in blood, earlier than those in pathological sections, and even earlier than the appearance of disease symptoms, suggesting that urine biomarkers can be applied for the early diagnosis of diseases. Since the urine proteome is susceptible to a variety of factors, such as diet, drug therapy, and daily activities, to make the experimental results more accurate, the key is to use a simple and controllable system. The genetic and environmental factors of animal models can be artificially controlled and can minimize the influence of unrelated factors, indicating that animal models are a very appropriate experimental method. For example, (1) Zhang Fansheng et al.^[3]^ found that the levels of 29 proteins changed in the urine before amyloid plaque deposition in the brains of transgenic mice with Alzheimer’s disease appeared, and 24 of them were reported to be associated with Alzheimer’s disease or used as markers. (2) Wu Jianqiang et al.^[4]^ demonstrated that in a Walker 256 subcutaneous tumor rat model, the levels of 10 proteins changed in the urine before the subcutaneous tumor was palpable. (3) Zhang Y et al.^[5]^ illustrated that in a chronic pancreatitis rat model, 15 differential proteins were identified in the urine before the pathology had changed at week 2, of which 5 were reported to be associated with pancreatitis; (4) Ni Yanying et al.^[6]^ showed that in a glioma rat model constructed by C6 cell injection into the brain, urinary protein levels changed before the appearance of pathological changes during imaging; (5) Zhang Fansheng et al.^[7]^ found that in a thioacetamide-induced liver fibrosis rat model, 40 differential proteins were identified in the urine before pathological changes, and 15 were reported to be associated with fibrosis. (6) Yin et al.^[8]^ found that urine glucose levels presented frequent disturbances before blood glucose increased in obese type 2 diabetic rats, which is important in indicating early diabetes. (7) Huang He et al.^[9]^ exposed rats to smoke from traditional cigarettes and screened biomarkers of chronic obstructive pulmonary disease (COPD) that have been reported only at two weeks of exposure. After comparative studies, it was found that urinary protein levels changed when tumor cells were subcutaneously injected^[4]^ and grew in different organs including liver^[7]^, bone^[10]^, lung^[11]^, and brain^[6]^, which indicated that urine had the potential to distinguish the growth of the same tumor cells in different organs. In terms of sample acquisition, urine acquisition is more noninvasive and easily available^[12]^, urine is a suitable source of biomarkers, and the construction of animal models is a very important means in urine proteomics research.

Currently, there are few studies on the effects of different odors on the rat urinary proteome. In a previous study, multiple isoforms of odor-binding proteins were identified in the urinary proteins of rats after smoking jasmine-flavored e-cigarettes, suggesting that the urinary proteome has the ability to reflect changes in the organism after smelling odors ^[13]^. So do rats produce different changes after smelling different odors? This is a very interesting scientific question. Therefore, in this paper, two kinds of odors, sesame oil and essential balm, were chosen to construct an animal model with 6-8 weeks old Wistar male rats, which is devoted to study the effect of smelling different odors on rat urine proteome (as in Figure 1), and provide a new method to explore the biological process of olfaction.

**Figure 1.**
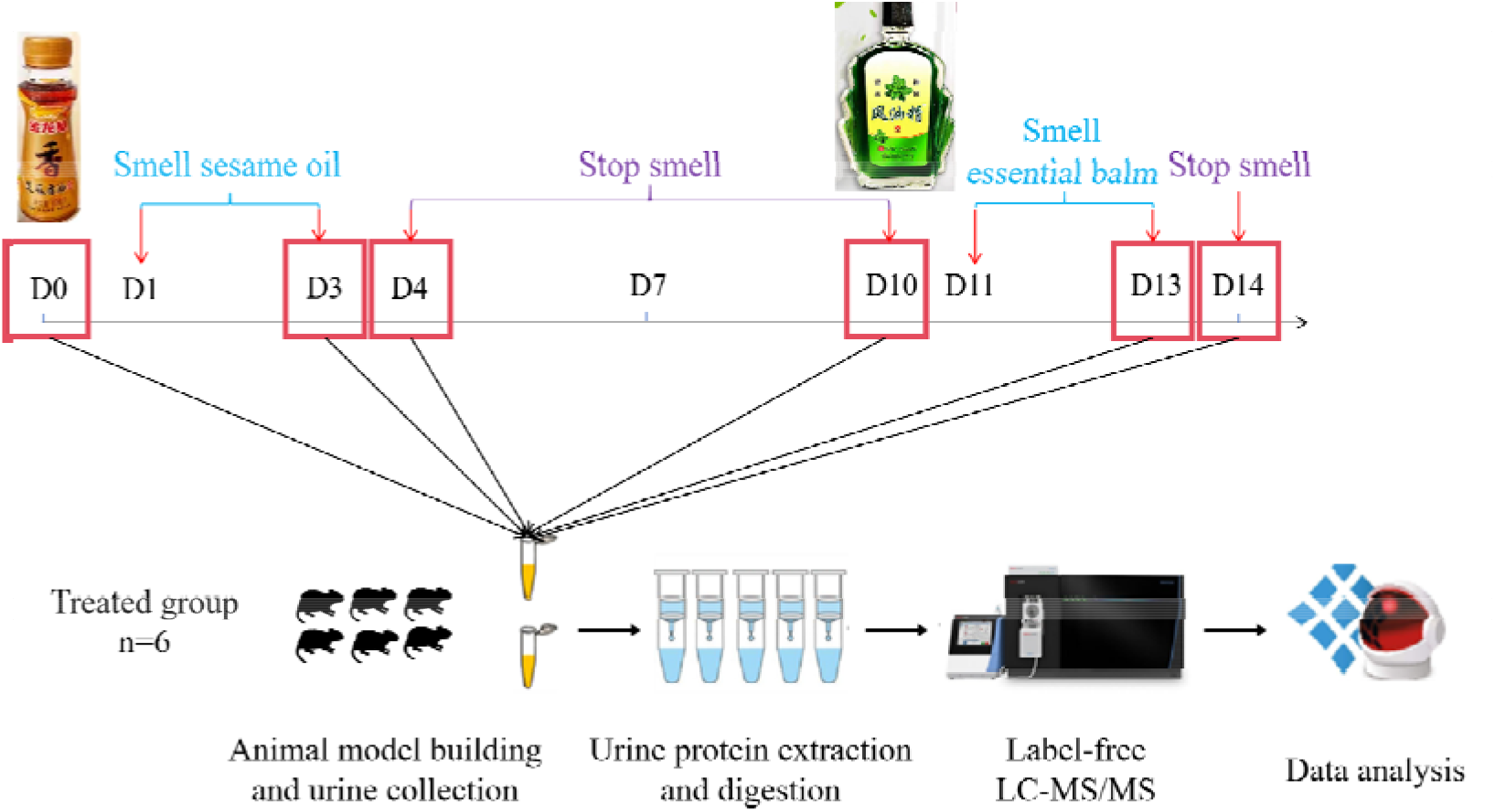
Workflow to investigate the effect of different odors on the rat urine proteome. Urine samples were collected on Days 0, 3, 4, 10, 13 and 14. After urine samples were collected and processed, the protein groups were identified using liquid chromatography coupled with tandem mass spectrometry (LC–MS/MS) to quantitatively analyze changes caused by different odors in rats.

## 2. Materials and Methods

### 2.1 Establishment of animal model

For this experiment, six healthy male Wistar rats (180-200 g) of 6-8 weeks old of SPF grade were selected and purchased from Beijing Beijing Vital River Laboratory Animal Technology Co. Ltd, with the animal license number of SYXK(Beijing)2021-0011.All rats were kept in a standard environment (room temperature (22±1)L, humidity 65%-70%). The experiment was started after keeping all rats in the new environment for three days, and all experimental operations followed the review and approval of the Ethics Committee of the School of Life Sciences, Beijing Normal University.

The animal model was established as follows: six rats of the experimental group smelled the sesame oil once at the same time period, and each time 15 mL of sesame oil was placed in a location inaccessible to the rats in the rat cage [36 cm (L) × 20 cm (W) × 28 cm (H)], and three rats of the experimental group were put into each cage for 1 hour under the condition of ensuring sufficient oxygenation for three consecutive days, and they were put back to their own cages after each smelling. After each smelling, they were put back to their own cages. After stopping the smelling of sesame oil for 1 week, the rats were made to smell essential balm once during the same period of time, and each time 15 mL of essential balm was placed in a location in the rat cage that was inaccessible to the rats, and 3 rats of the experimental group were put into each cage for 1 hour under the condition of ensuring sufficient oxygen content for 3 consecutive days of smelling, and then returned to their own cages after each smelling. Behavioral changes of the rats were observed during the experiment.

### 2.2 Urine collection

All rats were uniformly placed in metabolic cages to collect 12 h urine samples after three days of rearing in the new environment. On the 3rd day when the rats smelled the sesame oil and the 1st day when they stopped smelling the sesame oil (i.e., Day4), all the rats were placed in metabolic cages to collect urine samples for 12 h. The urine sample collection method for the essential balm group was the same as above. During the urine collection process, the rats were fasted and dehydrated, and the collected urine samples were stored in a −80°C refrigerator.

### 2.3 Treatment of the urine samples

Urine protein extraction and quantification: Rat urine samples collected at three time points were centrifuged at 12,000×g for 40 min at 4 °C, and the supernatants were transferred to new Eppendorf (EP) tubes. Three volumes of precooled absolute ethanol were added, homogeneously mixed and precipitated overnight at −20 °C. The following day, the mixture was centrifuged at 12,000×g for 30 min at 4 °C, and the supernatant was discarded. The protein pellet was resuspended in lysis solution (containing 8 mol/L urea, 2 mol/L thiourea, 25 mmol/L dithiothreitol, and 50 mmol/L Tris). The samples were centrifuged at 12,000×g for 30 min at 4 °C, and the supernatant was placed in a new EP tube. The protein concentration was measured using the Bradford assay.

Urinary protease cleavage: A 100 μg urine protein sample was added to the filter membrane (Pall, Port Washington, NY, USA) of a 10 kDa ultrafiltration tube and placed in an EP tube, and 25 mmol/L NH4HCO3 solution was added to make a total volume of 200 μL. Then, 20 mM dithiothreitol solution (dithiothreitol, DTT, Sigma) was added, and after vortex mixing, the metal bath was heated at 37 °C for 1 h and cooled to room temperature. Iodoacetamide (IAA, Sigma) was added at 50 mM, mixed well and allowed to react for 40 min at room temperature in the dark. Then, membrane washing was performed: □ 200 μL of UA solution (8 mol/L urea, 0.1 mol/L Tris-HCl, pH 8.5) was added and centrifuged twice at 14,000×g for 5 min at 18 °C; □ Loading: freshly treated samples were added and centrifuged at 14,000×g for 40 min at 18 °C; □ 200 μL of UA solution was added and centrifuged at 14,000×g for 40 min at 18 °C, repeated twice; □ 25 mmol/L NH4HCO3 solution was added and centrifuged at 14,000×g for 40 min at 18 °C, repeated twice; □ trypsin (Trypsin Gold, Promega, Trypchburg, WI, USA) was added at a ratio of 1:50 trypsin:protein for digestion and kept in a water bath overnight at 37 °C. The following day, peptides were collected by centrifugation at 13,000×g for 30 min at 4 °C, desalted through an HLB column (Waters, Milford, MA), dried using a vacuum dryer, and stored at −80 °C.

### 2.4 LC-MS/MS analysis

The digested samples were reconstituted with 0.1% formic acid, and the peptide sample concentrations were quantified using a BCA kit. They were then diluted to 0.5 μg/μL. Mixed peptide samples were prepared from 4 μL of each sample and separated using a high pH reversed-phase peptide separation kit (Thermo Fisher Scientific) according to the instructions. Ten effluents (fractions) were collected by centrifugation, dried using a vacuum dryer and reconstituted with 0.1% formic acid. iRT reagent (Biognosys, Switzerland) was added at a volume ratio of sample:iRT of 10:1 to calibrate the retention times of extracted peptide peaks. For analysis, 1 μg of each peptide from an individual sample was loaded onto a trap column and separated on a reverse-phase C18 column (50 μm×150 mm, 2 μm) using the EASY-nLC1200 HPLC system (Thermo Fisher Scientific, Waltham, MA). The elution for the analytical column lasted 90 min with a gradient of 5%-28% buffer B (0.1% formic acid in 80% acetonitrile; flow rate 0.3 μL/min). Peptides were analyzed with an Orbitrap Fusion Lumos Tribrid Mass Spectrometer (Thermo Fisher Scientific, MA).

To generate the spectrum library, 10 isolated fractions were subjected to mass spectrometry in data-dependent acquisition (DDA) mode. Mass spectrometry data were collected in high sensitivity mode. A complete mass spectrometric scan was obtained in the 350-1500 m/z range with a resolution set at 60,000. Individual samples were analyzed using Data Independent Acquisition (DIA) mode. DIA acquisition was performed using a DIA method with 36 windows. After every 10 samples, a single DIA analysis of the pooled peptides was performed as a quality control.

### 2.5 Database searching and label-free quantitation

Raw data collected from liquid chromatography□mass spectrometry were imported into Proteome Discoverer (version 2.1, Thermo Scientific) and the Swiss-Prot rat database (published in May 2019, containing 8086 sequences) for alignment, and iRT sequences were added to the rat database. Then, the search results were imported into Spectronaut Pulsar (Biognosys AG, Switzerland) for processing and analysis. Peptide abundances were calculated by summing the peak areas of the respective fragment ions in MS2. Protein intensities were summed from their respective peptide abundances to calculate total protein abundances.

### 2.6 Statistical analysis

Each sample was subjected to three technical replications and the mean was taken for statistical analysis. In this experiment, the samples from the experimental groups at different time periods were controlled before and after themselves, while the control group was set up to exclude differences in growth and development. The identified proteins were compared to screen for differential proteins. The loose screening conditions for differential proteins were: fold change between groups (FC, Fold change) ≥1.5 or ≤0.67, and P-value of two-tailed unpaired t-test <0.05. The strict screening conditions for differential proteins were: fold change between groups (FC, Fold change) ≥2 or ≤0.5, and P-value of two-tailed unpaired t-test <0.01. The differential proteins were screened by using the Wukong platform (https://www.omicsolution.org/wkomic/main/), which is the most popular platform in the world for screening differential proteins. proteins were functionally enriched using the Wukong platform, the Uniprot website (https://www.uniprot.org/) and DAVID database (https://david.ncifcrf.gov/) analysis. And the reported literature was searched in Pubmed database (https://pubmed.ncbi.nlm.nih.gov) so as to analyze the function of differential proteins.

## 3. Results

### 3.1 Characterization of e-cigarette smoking rats

In this experiment, the behavior of rats was observed when smelling sesame oil and essential balm. It was found that the rats were very excited and active when smelling the sesame oil and showed obvious willingness to consume it. On the other hand, when smelling the essential balm, the rats showed the behaviors of aversion, avoidance, and cowering in the corner to sleep.

### 3.2 Identification of the urinary proteome after smelling different odors in rats

After collecting urine samples from six 6-8-week-old male Wistar rats Day0, Day3, Day4, Day10, Day13, and Day14, peptides generated from enzymatic digestion of 36 urine samples were analyzed by LC-MS/MS tandem mass spectrometry. A total of 1145 proteins were identified (≥2 specific peptides, protein level FDR <1%).

### 3.3 Analysis of urinary protein changes in before and after comparisons within the odor-smelling group

#### 3.3.1 Urine proteome changes in rats before-after smelling sesame oil

Urinary proteins were compared at different time points after smelling sesame oil in rats. The criteria for screening differential proteins under relaxed conditions were FC ≥ 1.5 or ≤ 0.67, with a two-tailed unpaired t-test of P < 0.05. The results showed that 52 differential proteins could be identified when Day0 was compared with Day3 (Table S1). After comparing them with the differential proteins generated from the growth and development of rats of the same weekly age at an interval of 3 days (Table S2, Table S3), it was found that the two presented a low degree of overlap of differential proteins, with only 4 overlapping differential proteins out of the 52 differential proteins. It proves that the differential proteins identified this time are less likely to be produced by the growth and development of rats. 143 differential proteins could be identified when Day0 was compared with Day4 (Table S1), and included all the differential proteins screened by Day0 and Day3 (Figure 2). The screening criteria were FC ≥ 2 or ≤ 0.5 under stringent conditions with a two-tailed unpaired t-test of P < 0.01. The results showed that one differential protein, the Cd99 protein, could be identified on Day0 compared with Day3 with a Fold change value of 0.03 and a P-value of 8.80E-03. 17 differential proteins could be identified on Day0 compared with Day4( Table 1).

**Table 1.**
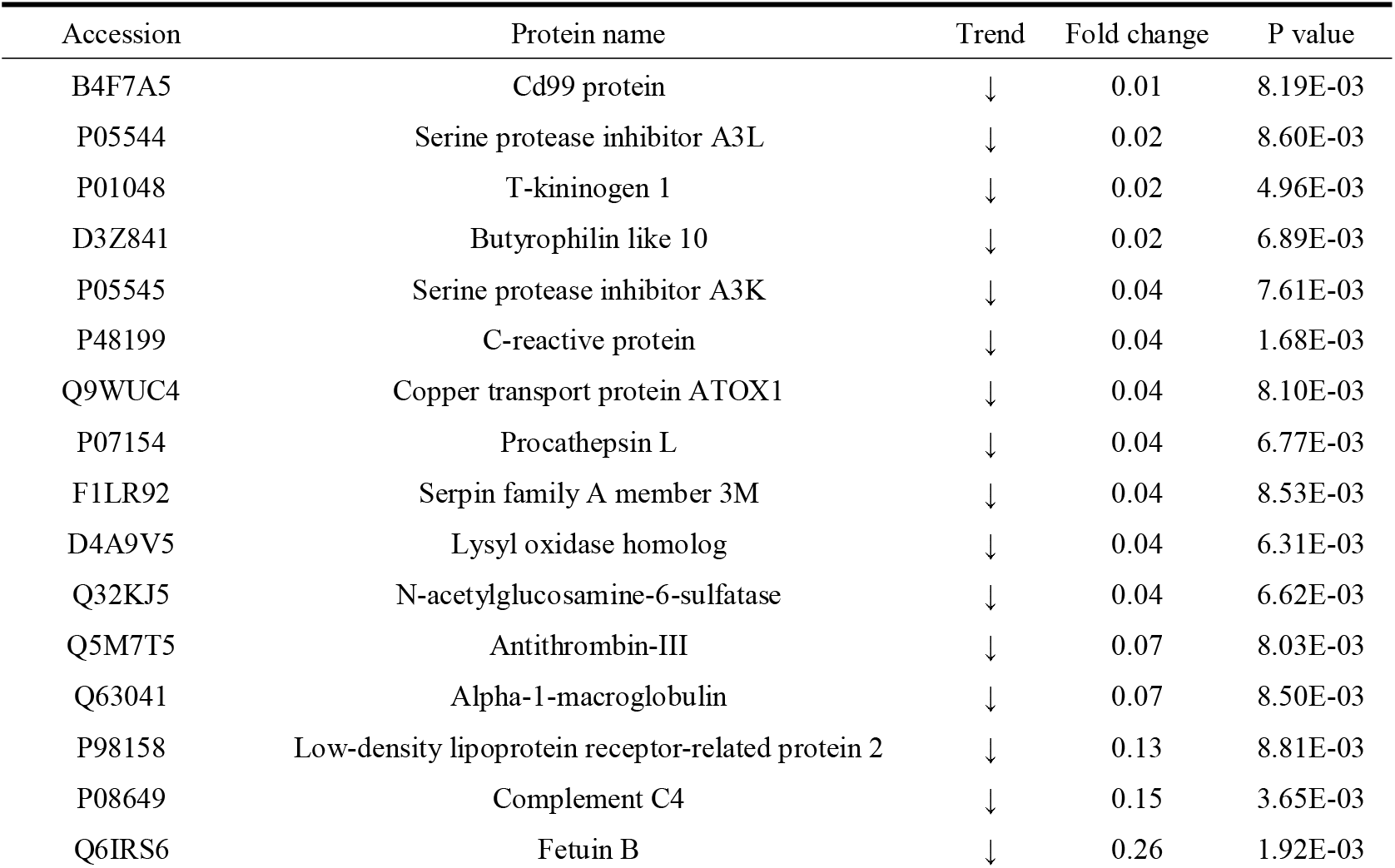
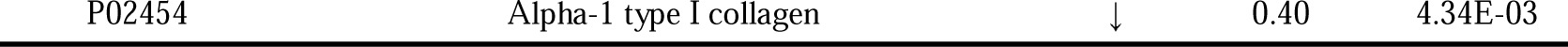
Differential proteins produced by Day0 vs Day4 rats smelling sesame oils(FC≥2 or ≤0.5, P<0.01)

| Accession | Protein name | Trend | Fold change | P value |
| --- | --- | --- | --- | --- |
| B4F7A5 | Cd99 protein | ↓ | 0.01 | 8.19E-03 |
| P05544 | Serine protease inhibitor A3L | ↓ | 0.02 | 8.60E-03 |
| P01048 | T-kininogen 1 | ↓ | 0.02 | 4.96E-03 |
| D3Z841 | Butyrophilin like 10 | ↓ | 0.02 | 6.89E-03 |
| P05545 | Serine protease inhibitor A3K | ↓ | 0.04 | 7.61E-03 |
| P48199 | C-reactive protein | ↓ | 0.04 | 1.68E-03 |
| Q9WUC4 | Copper transport protein ATOX1 | ↓ | 0.04 | 8.10E-03 |
| P07154 | Procathepsin L | ↓ | 0.04 | 6.77E-03 |
| F1LR92 | Serpin family A member 3M | ↓ | 0.04 | 8.53E-03 |
| D4A9V5 | Lysyl oxidase homolog | ↓ | 0.04 | 6.31E-03 |
| Q32KJ5 | N-acetylglucosamine-6-sulfatase | ↓ | 0.04 | 6.62E-03 |
| Q5M7T5 | Antithrombin-III | ↓ | 0.07 | 8.03E-03 |
| Q63041 | Alpha-1-macroglobulin | ↓ | 0.07 | 8.50E-03 |
| P98158 | Low-density lipoprotein receptor-related protein 2 | ↓ | 0.13 | 8.81E-03 |
| P08649 | Complement C4 | ↓ | 0.15 | 3.65E-03 |
| Q6IRS6 | Fetuin B | ↓ | 0.26 | 1.92E-03 |

**Figure 2.**
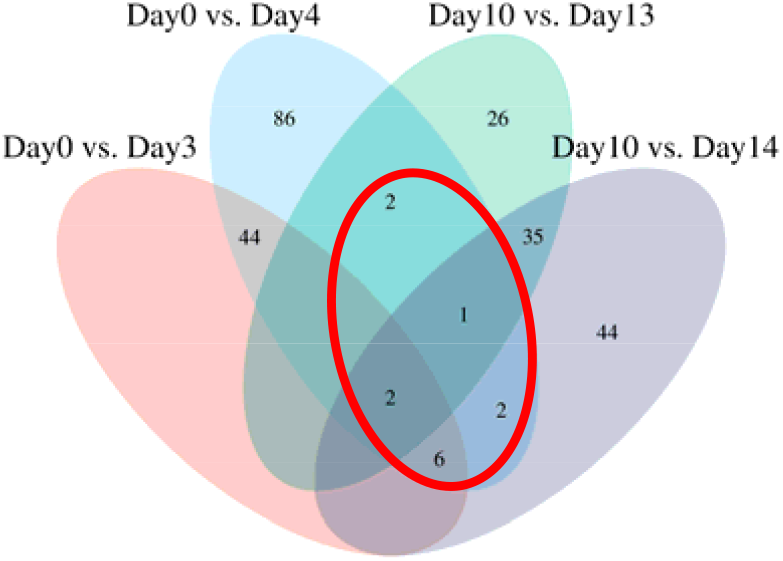
Differential protein Venn diagrams produced by smelling sesame oil and essential balm at different time points(FC≥1.5 or ≤0.67, P<0.05)

Among the differential proteins identified, several proteins have been reported to be associated with olfaction.Ekström I et al.^[14]^ in 2021 showed that serum C-reactive protein (CRP) levels were negatively correlated with olfactory recognition in aging. And Proft F et al.^[15]^ found that diminished sense of smell in patients with granulomatous polyangiitis (GPA) was significantly associated with elevated CRP values. Frizzled receptors (Frizzled) with conserved seven transmembrane helices are a class of atypical GPCRs and members of the Wnt/frizzled family mediate axon extension in olfactory sensory neurons^[16]^.Yue Y et al.^[17]^ further showed that Wnt-activated olfactory sheath cells, which are specialized glial cells located in the olfactory system, stimulate neural stem cells to proliferate and differentiate in a secretory manner. Strotmann J et al.^[18]^ used fluorescently labeled recombinant Odorant binding protein (OBP) loaded with odorant compounds and then applied to the noses of mice, and showed that the odorant compounds secreted by Low-density lipoprotein receptor-related protein 2 (LRP2, LRP2), which is a protein that is activated by Wnt, were secreted into the olfactory sheath cells. related protein 2 (LRP2) mediated internalization of the OBP/odorant complex into parasitic cells may be an important mechanism for the rapid and local removal of odorants.Gajera CR et al.^[19]^ demonstrated that a deficiency of LRP2 in adult mice resulted in impaired proliferation of neural precursor cells in the subventricular zone, which led to a decrease in the number of neuromasts arriving at the olfactory bulb. Diao WQ et al.^[20]^ demonstrated that fetuin B (Fetuin-B) is a plasma biomarker candidate that correlates with the severity of lung function in patients with chronic obstructive pulmonary disease (COPD). Apolipoprotein E (ApoE) (Table S1) was also identified in this study under relaxed conditions, and it has been well documented^[21][22]^ that ApoE is highly expressed in the central nervous system including the olfactory epithelium and olfactory bulb, and plays a key role in olfactory information processing in both the olfactory epithelium and olfactory bulb, which can affect the olfactory system in the early stages of Alzheimer’s disease. Struble RG et al.^[23]^ also showed that ApoE is associated with the ongoing degenerative and regenerative processes of olfactory neurogenesis.

Functional analysis of the differential proteins identified by Day0 compared with Day3 under relaxed conditions enriched 73 biological processes such as cellular response to cytochalasin B, negative regulation of endopeptidase activity, regulation of norepinephrine absorption, protein localization of adhesion junctions (Table S4) and 31 signaling pathways such as apoptosis, phagocytes, proteoglycans in cancer, and leukocyte transendothelial migration (Table S5). Functional analysis of the differential proteins identified by Day0 compared with Day4 under relaxed conditions enriched a total of 134 biological processes such as negative regulation of endopeptidase activity, proteolysis, cellular response to cytochalasin B, aging, acute phase response, and regulation of norepinephrine uptake (Table S6) and 34 signaling pathways such as proteoglycans, complement and coagulation cascades, and leukocyte transendothelial migration in cancer (Table S7).

#### 3.3.2 Urine proteome changes in rats before-after smelling essential balm

Urinary proteins at different time points after smelling essential balm in rats were compared. The criteria for screening differential proteins under relaxed conditions were FC ≥ 1.5 or ≤ 0.67, two-tailed unpaired t-test P < 0.05. The results showed that 66 differential proteins could be identified on Day10 compared with Day13 (Table S8). After comparing them with the differential proteins generated from the growth and development of rats of the same weekly age at an interval of 3 days (Table S2, Table S3), it was found that the two presented a low degree of overlap of differential proteins, with only 2 differential proteins out of the 66 differential proteins overlapping. It proves that the differential proteins identified this time are less likely to be produced by rat growth and development. 90 differential proteins could be identified when Day10 was compared with Day14 (Table S8). The screening criteria were FC ≥ 2 or ≤ 0.5 under stringent conditions with two-tailed unpaired t-test P < 0.01. The results showed that 14 differential proteins could be identified when Day10 was compared with Day13 (Table 2). 20 differential proteins could be identified when Day10 was compared with Day14 (Table 2), of which 10 differential proteins were duplicated by Day10 compared with Day13.

**Table 2.**
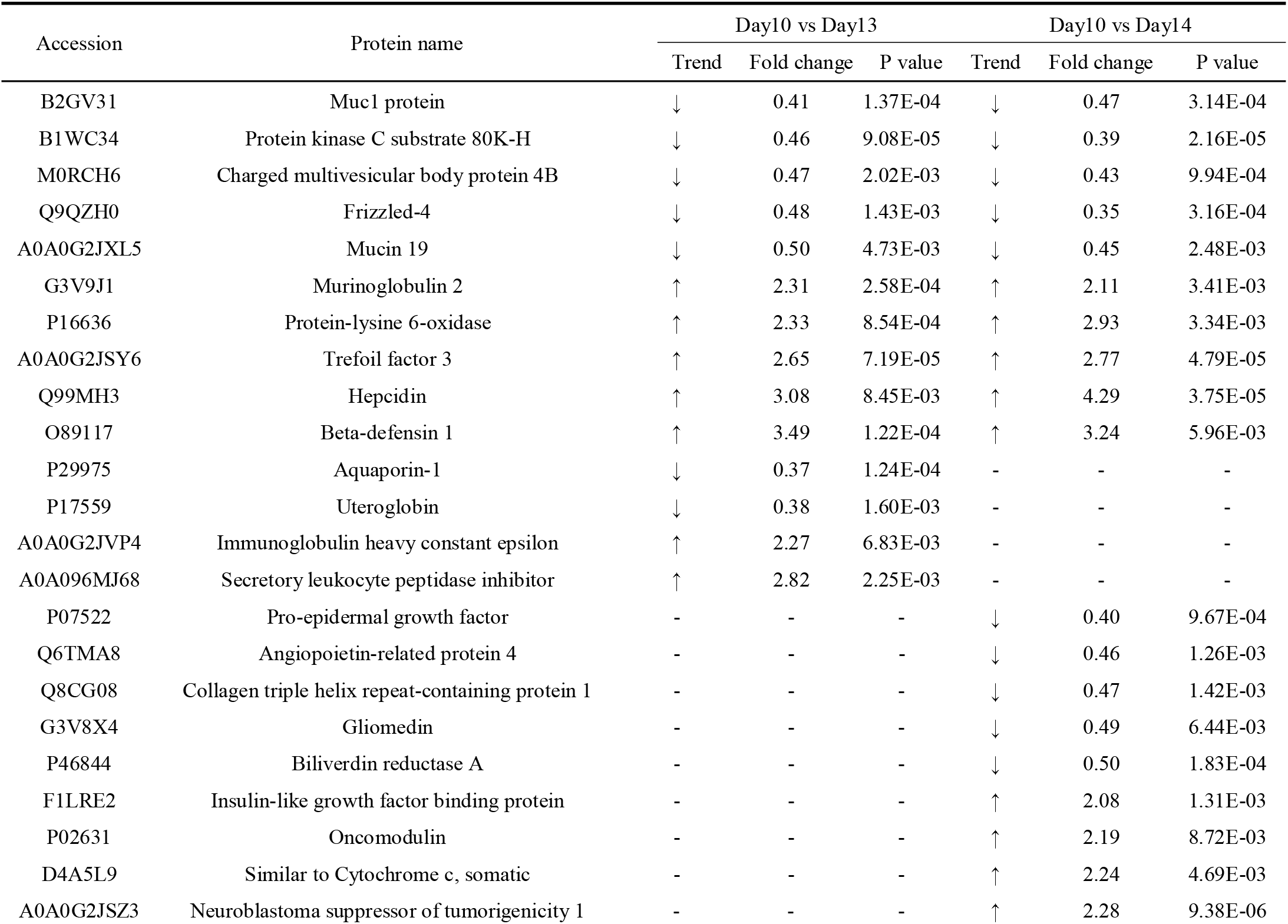
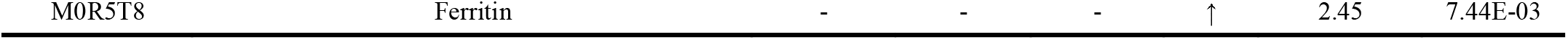
Differential proteins produced by smelling essential balm in rats (FC≥2 or ≤0.5, P<0.01)

| Accession | Protein name | Day10 vs Day13 |  |  | Day10 vs Day14 |  |  |
| --- | --- | --- | --- | --- | --- | --- | --- |
|  |  | Trend | Fold change | P value | Trend | Fold change | P value |
| B2GV31 | Muc1 protein | ↓ | 0.41 | 1.37E-04 | ↓ | 0.47 | 3.14E-04 |
| B1WC34 | Protein kinase C substrate 80K-H | ↓ | 0.46 | 9.08E-05 | ↓ | 0.39 | 2.16E-05 |
| M0RCH6 | Charged multivesicular body protein 4B | ↓ | 0.47 | 2.02E-03 | ↓ | 0.43 | 9.94E-04 |
| Q9QZH0 | Frizzled-4 | ↓ | 0.48 | 1.43E-03 | ↓ | 0.35 | 3.16E-04 |
| A0A0G2JXL5 | Mucin 19 | ↓ | 0.50 | 4.73E-03 | ↓ | 0.45 | 2.48E-03 |
| G3V9J1 | Murinoglobulin 2 | ↑ | 2.31 | 2.58E-04 | ↑ | 2.11 | 3.41E-03 |
| P16636 | Protein-lysine 6-oxidase | ↑ | 2.33 | 8.54E-04 | ↑ | 2.93 | 3.34E-03 |
| A0A0G2JSY6 | Trefoil factor 3 | ↑ | 2.65 | 7.19E-05 | ↑ | 2.77 | 4.79E-05 |
| Q99MH3 | Hepcidin | ↑ | 3.08 | 8.45E-03 | ↑ | 4.29 | 3.75E-05 |
| O89117 | Beta-defensin 1 | ↑ | 3.49 | 1.22E-04 | ↑ | 3.24 | 5.96E-03 |
| P29975 | Aquaporin-1 | ↓ | 0.37 | 1.24E-04 | - | - | - |
| P17559 | Uteroglobin | ↓ | 0.38 | 1.60E-03 | - | - | - |
| A0A0G2JVP4 | Immunoglobulin heavy constant epsilon | ↑ | 2.27 | 6.83E-03 | - | - | - |
| A0A096MJ68 | Secretory leukocyte peptidase inhibitor | ↑ | 2.82 | 2.25E-03 | - | - | - |
| P07522 | Pro-epidermal growth factor | - | - | - | ↓ | 0.40 | 9.67E-04 |
| Q6TMA8 | Angiopoietin-related protein 4 | - | - | - | ↓ | 0.46 | 1.26E-03 |
| Q8CG08 | Collagen triple helix repeat-containing protein 1 | - | - | - | ↓ | 0.47 | 1.42E-03 |
| G3V8X4 | Gliomedin | - | - | - | ↓ | 0.49 | 6.44E-03 |
| P46844 | Biliverdin reductase A | - | - | - | ↓ | 0.50 | 1.83E-04 |
| F1LRE2 | Insulin-like growth factor binding protein | - | - | - | ↑ | 2.08 | 1.31E-03 |
| P02631 | Oncomodulin | - | - | - | ↑ | 2.19 | 8.72E-03 |
| D4A5L9 | Similar to Cytochrome c, somatic | - | - | - | ↑ | 2.24 | 4.69E-03 |
| A0A0G2JSZ3 | Neuroblastoma suppressor of tumorigenicity 1 | - | - | - | ↑ | 2.28 | 9.38E-06 |

Under stringent conditions, differential proteins such as mucin 1 (Muc1), protein kinase C substrate 80K-H, mucin 19, cloverleaf factor 3, and β-defensin 1 (BD-1) were identified in Day10 compared with Day13 and Day10 compared with Day14 (Table 2).Of the 21 known mucin isozymes, mucins 1, 4, 5AC, and 8 are found in human sinus epithelium and mucins 1, 5B, and 8 in sinus glands^[24][25][26][27]^.Some mucins, including Muc1, have transmembrane peptides that bind to cell membranes, whereas others are secreted completely^[28]^, suggesting that mucin 1 may protect extremely sensitive structures such as olfactory neurons^[29]^.Bruch RC et al. showed that protein kinase C (PKC), together with G protein-coupled receptor kinase, mediates signal termination through phosphorylation of odorant receptors and possibly other substrates and is involved in termination and desensitization of olfactory signals^[30][31]^. Li J et al.^[32]^ showed that TFF3 reversed depression-like behavior in olfactory bulbectomy rats by activating BDNF-ERK-CREB signaling.Baines KJ et al.^[33]^ showed that elevated β-defensin-1 protein is a feature of COPD and severe asthma, and dysregulated production of β-defensin-1 in epithelial cells of COPD patients demonstrates that it may be an effective biomarker and potential therapeutic target for COPD.

Differential proteins such as aquaporin-1 (AQP-1) were identified in Day10 compared with Day13 under stringent conditions (Table 2) and Day10 compared with Day14 under relaxed conditions (Table S8).It has been shown that AQP-1 is expressed in olfactory ensheathing glial cells, vascular endothelial cells of olfactory and respiratory mucosa, and surrounding connective tissue cells^[34][35]^.However, its expression is not specific, and the association with olfaction is unknown.Differential proteins such as gliomedin and insulin-like growth factor binding protein (IGFBP) were identified in Day10 compared with Day14 under stringent conditions (Table 2) and Day10 compared with Day13 under relaxed conditions (Table S8).Gliomedin is a member of the olfactomedin protein family and is required for molecular assembly of Ranvier developmental nodes in the peripheral nervous system^[36][37]^.IGFBP-2 binds to proteoglycans in rat olfactory bulb membranes and regulates chitosan-mediated neuronal differentiation of human olfactory receptors^[38][39]^.Pro-epidermal growth factor (EGF) identified on Day10 compared with Day14 under relaxed conditions (Table S8), Farbman AI et al.^[40]^ showed that EGF may be involved in mitotic regulation of olfactory epithelial cells, while Ohta Y et al.^[41]^ compared the number of regions where epidermal growth factor receptors were found in the basal layer of the olfactory epithelium between aged mice and adult mice and concluded that reduction of epidermal growth factor receptors in the olfactory epithelium inhibits cell proliferation and thus leads to olfactory epithelial atrophy.

Functional analysis of the differential proteins identified on Day10 compared with Day13 under relaxed conditions enriched a total of 29 biological processes including intermediate filament tissue, wound healing, and response to nutrient levels (Table S9) and three signaling pathways: S. aureus infection, ECM interaction with receptors, and TGF-β signaling pathway (Table S10). Functional analysis of the differential proteins identified on Day10 compared with Day14 under relaxed conditions enriched a total of 142 biological processes such as positive regulation of cell migration, response to xenobiotics (Table S11), and eight signaling pathways such as lesion adhesion, fluid shear stress, and proteoglycans in atherosclerosis and cancer (Table S12).

### 3.4 Analysis of urine protein changes between groups after smelling

Venn diagrams of the differential proteins produced by the comparison of the four groups under relaxed conditions were plotted (Figure 2), and the results showed that only 13 differential proteins were common to the sesame oil group and the essential balm group (Table 3), and most of the differential proteins were concentrated in the groups with different odors. It shows that the consistency of the effect of different odors on the production of rat urine proteome is small.

**Table 3.**
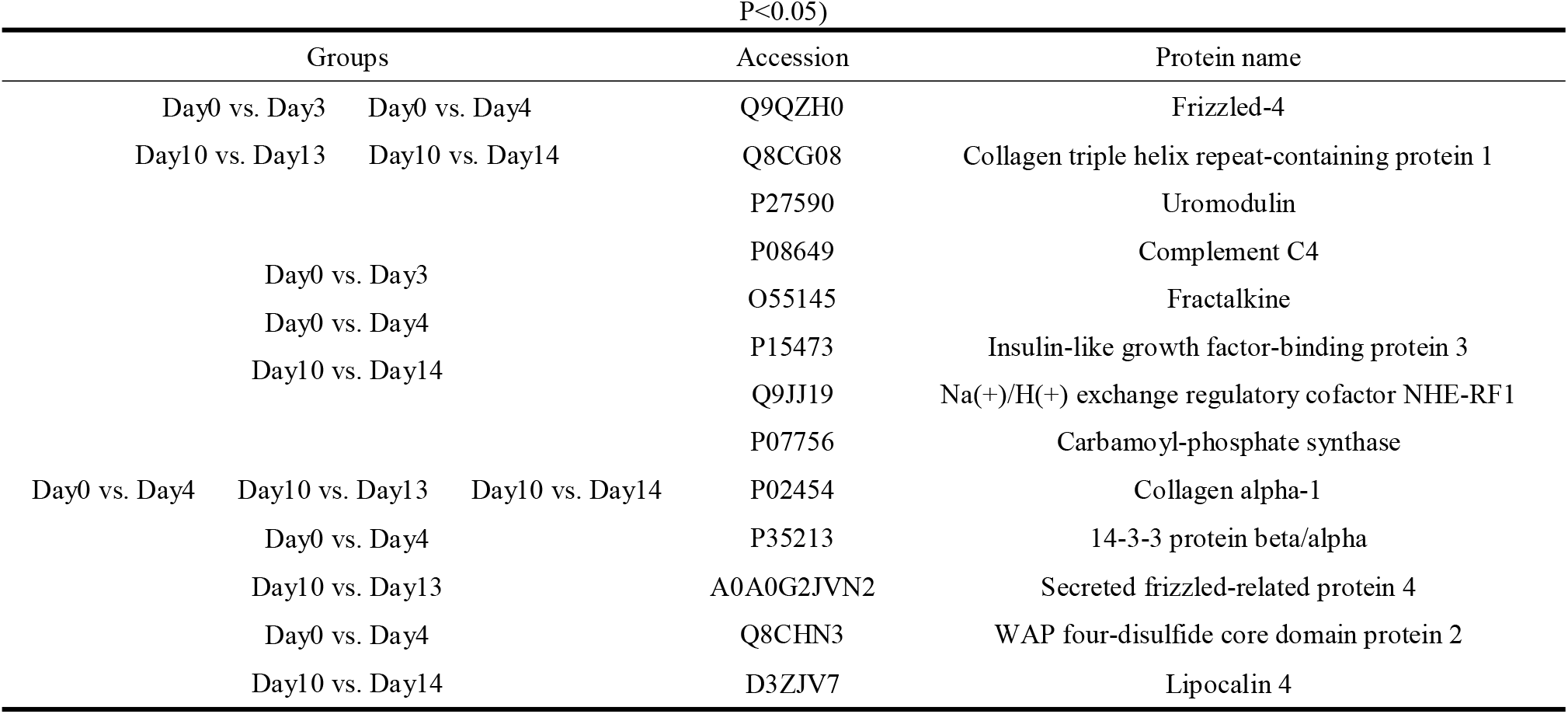
Differential proteins co-identified in the sesame oil group and the essential balm group(FC≥1.5 or ≤0.67, 或<0.05)

Very interestingly, we found serine-related proteins several times in the sesame oil group and the essential balm group (Table 4). It has been shown that overexpression of the serine protease inhibitor Spi2 in vitro produced a secretory signal for olfactory neuron death, suggesting that the serine protease inhibitor Spi2 may mediate olfactory neuron apoptosis^[42]^. In contrast, the calcium/calmodulin-dependent serine protein kinase CASK is present in olfactory cilia and may play a role in odor transmission^[43]^. No study has yet reported the relationship between the serine-related proteins identified in this study and olfactory production, which may provide clues for further investigation of the mechanism of olfactory production.

**Table 4.**
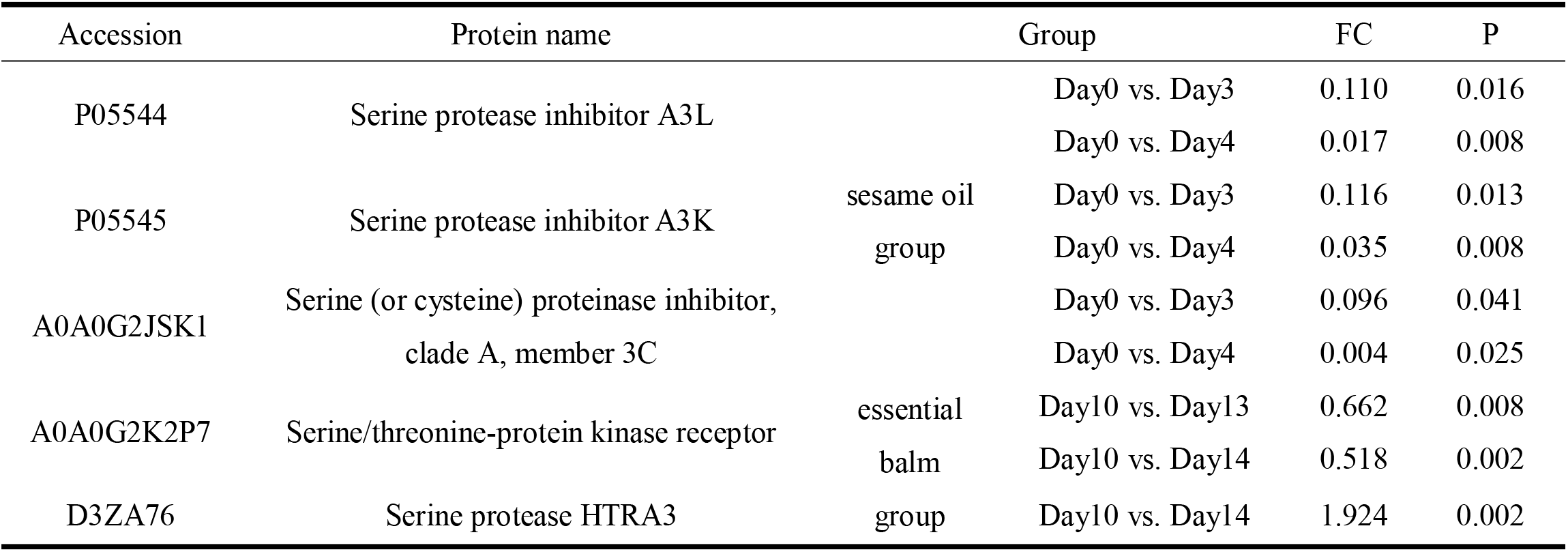
Proteins related to serine identified in the sesame oil and essential balm groups(FC≥1.5 or ≤0.67, P<0.05)

**Table 5.**
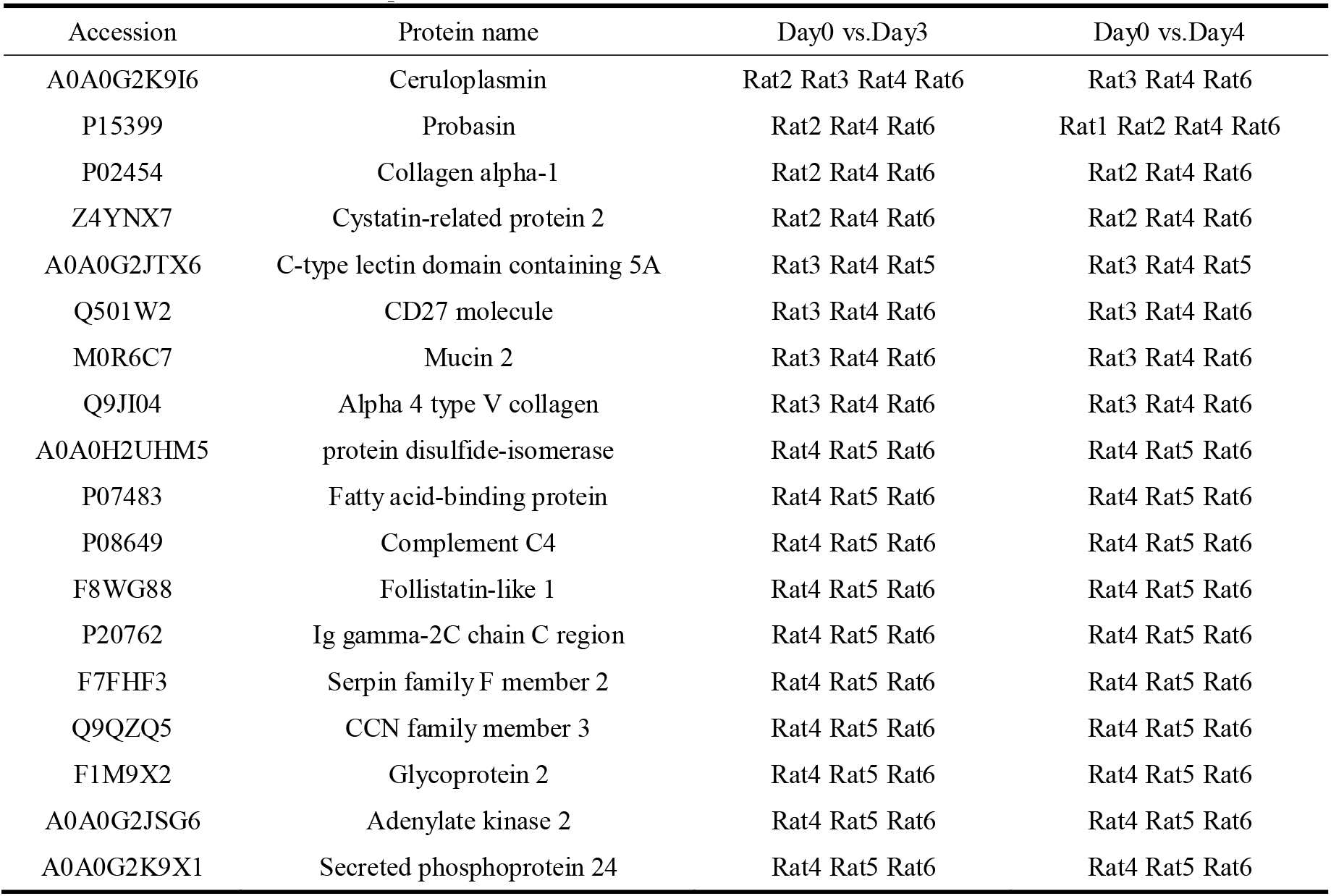
Differential proteins co-identified by three or more rats in the sesame oil group before and after comparison of six rats themselves(FC≥1.5 or ≤0.67, P<0.05)

| Accession | Protein name | Day0 vs.Day3 | Day0 vs.Day4 |
| --- | --- | --- | --- |
| A0A0G2K9I6 | Ceruloplasmin | Rat2 Rat3 Rat4 Rat6 | Rat3 Rat4 Rat6 |
| P15399 | Probasin | Rat2 Rat4 Rat6 | Rat1 Rat2 Rat4 Rat6 |
| P02454 | Collagen alpha-1 | Rat2 Rat4 Rat6 | Rat2 Rat4 Rat6 |
| Z4YNX7 | Cystatin-related protein 2 | Rat2 Rat4 Rat6 | Rat2 Rat4 Rat6 |
| A0A0G2JTX6 | C-type lectin domain containing 5A | Rat3 Rat4 Rat5 | Rat3 Rat4 Rat5 |
| Q501W2 | CD27 molecule | Rat3 Rat4 Rat6 | Rat3 Rat4 Rat6 |
| M0R6C7 | Mucin 2 | Rat3 Rat4 Rat6 | Rat3 Rat4 Rat6 |
| Q9JI04 | Alpha 4 type V collagen | Rat3 Rat4 Rat6 | Rat3 Rat4 Rat6 |
| A0A0H2UHM5 | protein disulfide-isomerase | Rat4 Rat5 Rat6 | Rat4 Rat5 Rat6 |
| P07483 | Fatty acid-binding protein | Rat4 Rat5 Rat6 | Rat4 Rat5 Rat6 |
| P08649 | Complement C4 | Rat4 Rat5 Rat6 | Rat4 Rat5 Rat6 |
| F8WG88 | Follistatin-like 1 | Rat4 Rat5 Rat6 | Rat4 Rat5 Rat6 |
| P20762 | Ig gamma-2C chain C region | Rat4 Rat5 Rat6 | Rat4 Rat5 Rat6 |
| F7FHF3 | Serpin family F member 2 | Rat4 Rat5 Rat6 | Rat4 Rat5 Rat6 |
| Q9QZQ5 | CCN family member 3 | Rat4 Rat5 Rat6 | Rat4 Rat5 Rat6 |
| F1M9X2 | Glycoprotein 2 | Rat4 Rat5 Rat6 | Rat4 Rat5 Rat6 |
| A0A0G2JSG6 | Adenylate kinase 2 | Rat4 Rat5 Rat6 | Rat4 Rat5 Rat6 |
| A0A0G2K9X1 | Secreted phosphoprotein 24 | Rat4 Rat5 Rat6 | Rat4 Rat5 Rat6 |

### 3.5 Analysis of urinary protein changes in individual rats before-after smelling

#### 3.5.1 Analysis of urinary protein changes in individual rats before-after smelling sesame oil

Venn diagrams were produced by comparing the differential proteins produced before and after Day0 and Day3 itself of the six experimental groups of rats smelling sesame oil under relaxed conditions (Figure 3A). The results showed that there were no differential proteins common to all five, and the number of differential proteins identified in rats 1 and 6 showed a large gap under the same screening conditions, indicating that the changes in the urinary proteome of the rats after smelling of sesame oil were characterized by strong individual variation. This was also illustrated by the Venn diagram presented on Day0 compared with Day4 (Figure 3B).

**Fig 3.**
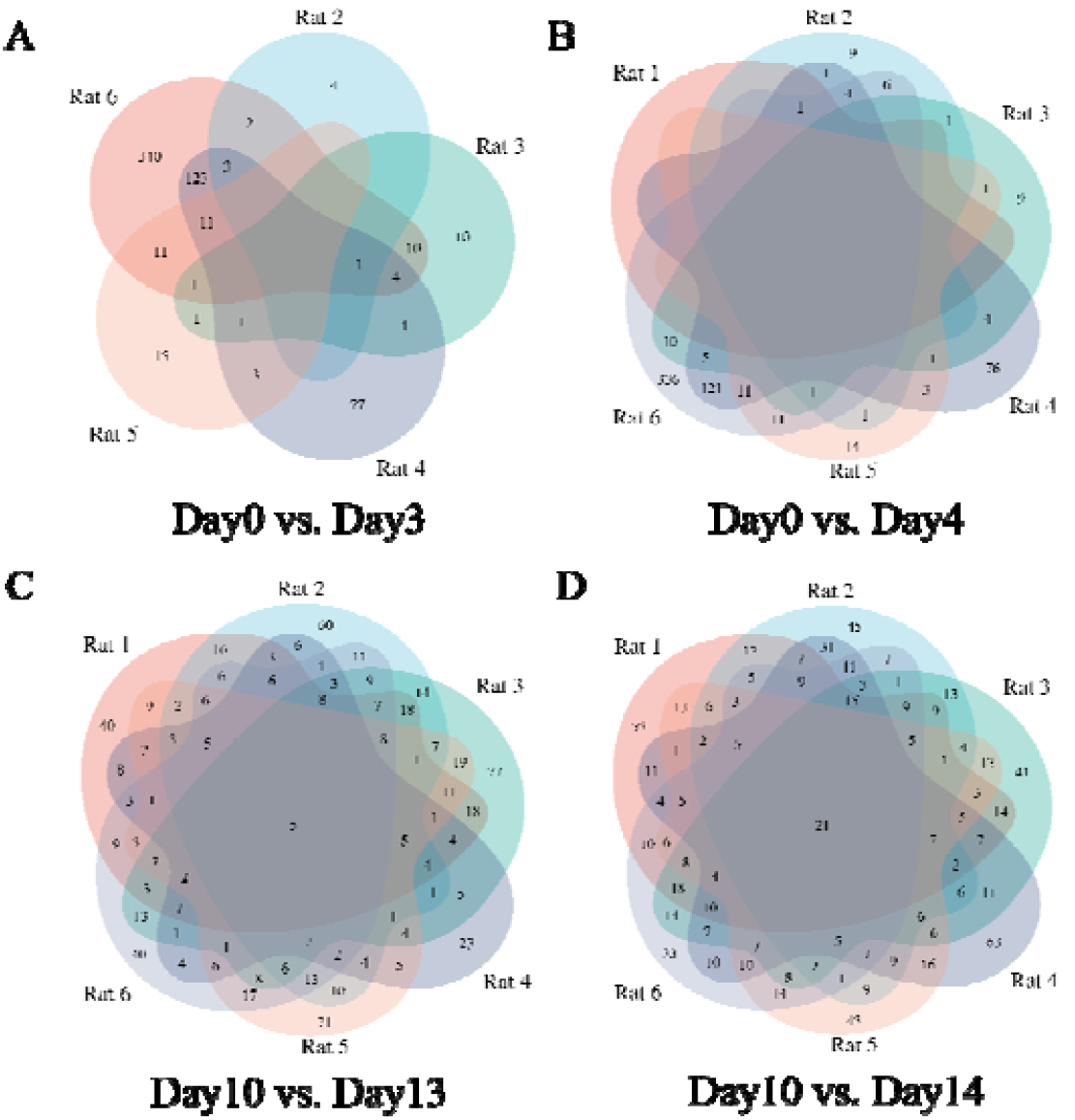
Venn diagram of the differential proteins produced by the before-after comparisons of the six rats themselves in the sesame oil group and the essential balm group at different time points.(FC≥1.5 or ≤0.67, P<0.05)Day0 vs. Day3. (B) Day0 vs. Day4. (C) Day10 vs. Day13. (D) Day10 vs. Day14.

#### 3.5.2 Analysis of urinary protein changes in individual rats before-after smelling essential balm

Under relaxed conditions, Venn diagrams were made of the differential proteins produced by comparing Day10 and Day13 of essential balm smelling from six experimental group rats before and after themselves (Figure 3C). The results showed that 18 differential proteins were common to 6 rats after smelling essential balm(Table 6), while 63 differential proteins were common to 5 and more rats (Table S13). Compared with the strong individual variation in the sesame oil group, the essential balm group showed more consistent changes as a whole. This is also illustrated by the Venn diagrams presented on Day10 compared to Day14 (Figure 3D). Among them, differential proteins such as clover factor 3, hepcidin, and mucin were similarly identified in group analysis (Table 2, Table 6). We also identified three isoforms of phosphatidylinositol proteoglycans (Glypican) (Glypican1, 3, 4) in multiple rats. Among them, studies by Saroja SR et al.^[44]^ demonstrated that Glypican4 is a binding partner of Apoe4, and Apoe4-induced tau hyperphosphorylation is directly mediated by Glypican4. Apoe4, which we identified at the time of comparative analysis of sesame oil composition groups, plays a key role in olfactory information processing in the olfactory epithelium and olfactory bulb and can influence the olfactory system early in Alzheimer ‘s disease^[21][22]^. While the relationship between Glypican4 and olfaction has not yet been reported, this study may provide clues for research in this direction.

**Table 6.**
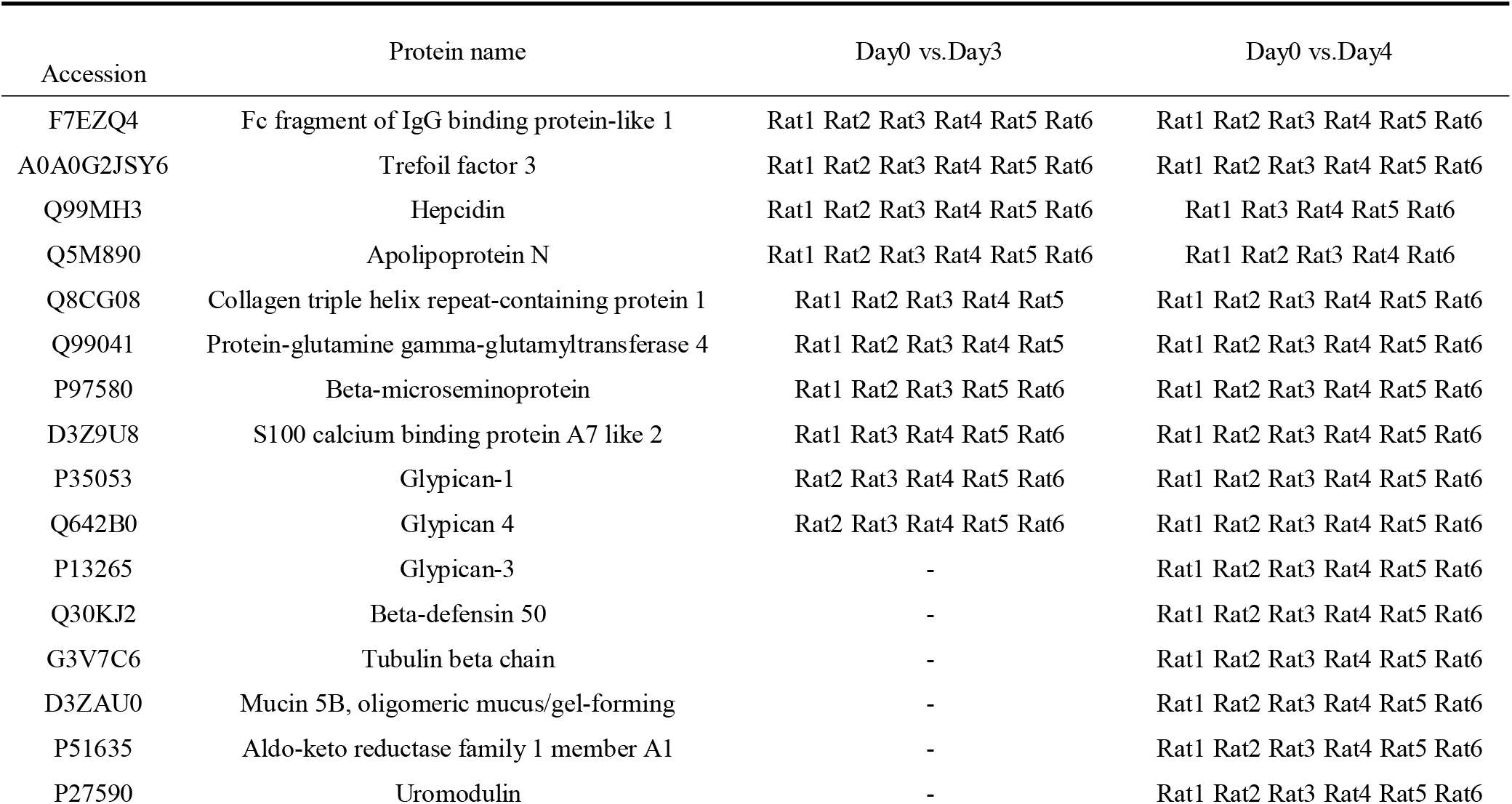
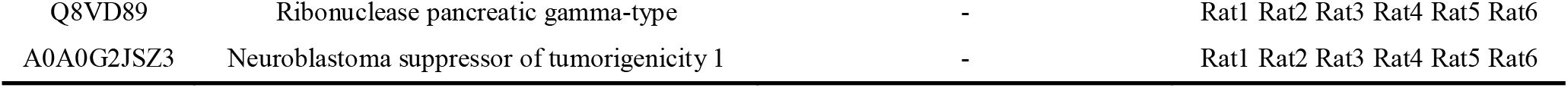
Differential proteins co-identified in 6 and more rats in the essential balm group on different days (FC ≥1.5 or ≤0.67, P<0.05)

It is worth mentioning that Odorant binding protein 2A (OBP2A) was identified in a total of 4 rats in the Day10 vs. Day13 group, and 3 of them showed a down-regulation trend. OBP2A was identified in a total of 3 rats in the Day10 and Day14 groups, and 2 of them showed a down-regulation trend. This protein was also identified in the rat e-cigarette model, and the changes in the expression of this protein showed strong consistency among the six rats^[13]^.

### 3.6 Randomization test

To determine the likelihood that the identified differential proteins were randomly generated, the total proteins identified from the six samples collected on Day0 and the six samples collected on Day3 were verified in randomized groups (FC ≥ 1.5 or ≤ 0.67, P<0.05), and there were a total of 462 different combinatorial scenarios that yielded an average of 9.79 differential proteins, suggesting that at least 81.17% of differential proteins were not randomly generated (Table 7). Randomized group validation of the total proteins identified in the six samples collected on Day0 and the six samples collected on Day4 (FC≥1.5 or ≤0.67, P<0.05) produced an average of 7.42 differential proteins, indicating that at least 94.88% of the differential proteins were not randomly generated (Table 7); randomized group validation with more stringent conditions (FC≥2 or ≤0.5, P<0.01) for randomized group validation produced an average of 0.16 differential proteins, indicating that at least 99.06% of the differential proteins were not randomly generated (Table 7). Randomized group validation of total proteins identified in the 6 samples collected on Day10 and the 6 samples collected on Day13 (FC≥1.5 or ≤0.67, P<0.05) produced an average of 19.91 differential proteins, suggesting that at least 69.83% of the differential proteins were not randomly generated (Table 7); randomized group validation of total proteins identified under more stringent conditions (FC≥2 or ≤0.5, P< 0.01) randomized group validation produced a mean of 1.41 differential proteins, indicating that at least 89.93% of the differential proteins were not randomly generated (Table 7). Randomized group validation of total proteins identified in the 6 samples collected on Day10 and the 6 samples collected on Day14 (FC ≥1.5 or ≤0.67, P<0.05) yielded an average of 21.12 differential proteins, indicating that at least 76.53% of the differential proteins were not randomly generated (Table 7); randomized group validation of total proteins identified under more stringent conditions (FC ≥2 or ≤0.5, P< 0.01) for randomized group validation produced a mean of 1.49 differential proteins, indicating that at least 92.55% of the differential proteins were not randomly generated (Table 7).

**Table 7.**
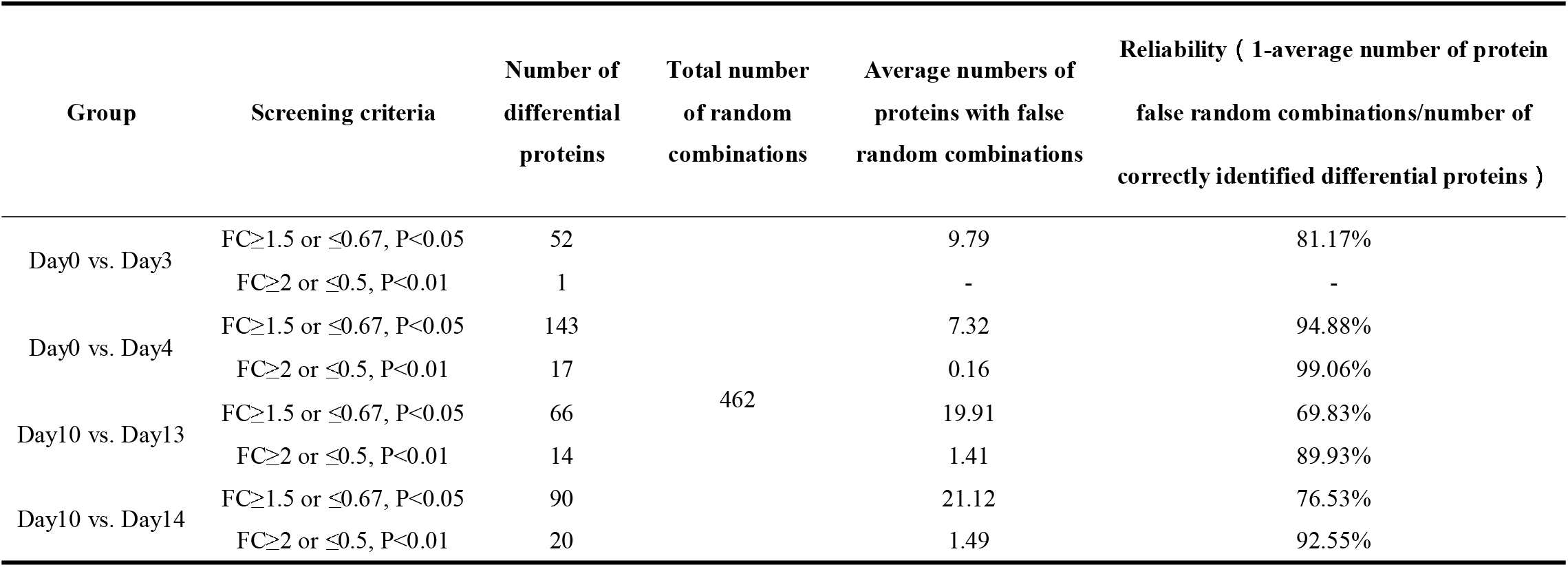
Randomization results.

| Group | Screening criteria | Number of differential proteins | Total number of random combinations | Average numbers of proteins with false random combinations | Reliability ( 1-average number of protein false random combinations/number of correctly identified differential proteins ) |
| --- | --- | --- | --- | --- | --- |
| Day0 vs. Day3 | $FC \geq 1.5$ or $\leq 0.67$, $P < 0.05$ | 52 | 462 | 9.79 | 81.17% |
| | $FC \geq 2$ or $\leq 0.5$, $P < 0.01$ | 1 | | - | - |
| Day0 vs. Day4 | $FC \geq 1.5$ or $\leq 0.67$, $P < 0.05$ | 143 | | 7.32 | 94.88% |
| | $FC \geq 2$ or $\leq 0.5$, $P < 0.01$ | 17 | | 0.16 | 99.06% |
| Day10 vs. Day13 | $FC \geq 1.5$ or $\leq 0.67$, $P < 0.05$ | 66 | | 19.91 | 69.83% |
| | $FC \geq 2$ or $\leq 0.5$, $P < 0.01$ | 14 | | 1.41 | 89.93% |
| Day10 vs. Day14 | $FC \geq 1.5$ or $\leq 0.67$, $P < 0.05$ | 90 | | 21.12 | 76.53% |
| | $FC \geq 2$ or $\leq 0.5$, $P < 0.01$ | 20 | | 1.49 | 92.55% |

## 4. Conclusion

The present study shows the effect that odor can produce on rat urine proteome. And different odors, such as sesame oil and essential balm, produce different effects on the rat urine proteome. This provides a new method to explore the biological process of olfaction.

