## Supplemental table1-13 for "Effect of different odors on the rat urine proteome"

Table S1 Differential proteins produced by sniffing aromatic oils in rats (FC≥1.5 or ≤0.67, P<0.05)

| Accession | Protein names | Day0 vs Day3 | | | | Day0 vs Day4 | | |
| --- | --- | --- | --- | --- | --- | --- | --- | --- |
|  |  | Trend | Fold change | | P value | Trend | Fold change | P value |
| P08721 | Osteopontin | ↓ | 0.01 | 1.89E-02 | | ↓ | 0.002 | 1.85E-02 |
| O55145 | Fractalkine | ↓ | 0.02 | 4.64E-02 | | ↓ | 0.001 | 4.23E-02 |
| B4F7A5 | Cd99 protein | ↓ | 0.03 | 8.80E-03 | | ↓ | 0.01 | 8.19E-03 |
| A0A0G2JSP1 | Uromodulin | ↓ | 0.03 | 1.46E-02 | | ↓ | 0.0003 | 1.26E-02 |
| Q64319 | Amino acid transporter heavy chain SLC3A1 | ↓ | 0.03 | 3.89E-02 | | ↓ | 0.01 | 3.53E-02 |
| E9PU07 | Ribonuclease A f4 | ↓ | 0.03 | 4.63E-02 | | ↓ | 0.001 | 4.09E-02 |
| Q6IE52 | Murinoglobulin-2 | ↓ | 0.03 | 1.79E-02 | | ↓ | 0.02 | 1.70E-02 |
| Q9JJ19 | Na(+)/H(+) exchange regulatory cofactor NHE-RF1 | ↓ | 0.03 | 1.52E-02 | | ↓ | 0.02 | 1.37E-02 |
| P27590 | Uromodulin | ↓ | 0.04 | 2.32E-02 | | ↓ | 0.002 | 1.92E-02 |
| G3V8G5 | Golgi apparatus protein 1 | ↓ | 0.06 | 2.69E-02 | | ↓ | 0.03 | 2.37E-02 |
| Q4QQW8 | Putative phospholipase B-like 2 | ↓ | 0.06 | 4.39E-02 | | ↓ | 0.03 | 3.73E-02 |
| P07632 | Superoxide dismutase [Cu-Zn] | ↓ | 0.07 | 2.71E-02 | | ↓ | 0.01 | 1.97E-02 |
| G3V8M6 | Folate receptor alpha | ↓ | 0.07 | 2.32E-02 | | ↓ | 0.01 | 1.66E-02 |
| P07861 | Neprilysin | ↓ | 0.07 | 2.70E-02 | | ↓ | 0.05 | 2.40E-02 |
| P15473 | Insulin-like growth factor-binding protein 3 | ↓ | 0.07 | 1.89E-02 | | ↓ | 0.04 | 1.52E-02 |
| A0A0A0MXX5 | 3-hydroxybutyrate dehydrogenase 2 | ↓ | 0.08 | 4.15E-02 | | ↓ | 0.001 | 2.84E-02 |
| Q03626 | Murinoglobulin-1 | ↓ | 0.08 | 1.81E-02 | | ↓ | 0.02 | 1.31E-02 |
| P07314 | Glutathione hydrolase 1 proenzyme | ↓ | 0.09 | 2.14E-02 | | ↓ | 0.02 | 1.36E-02 |
| F1LNY3 | Neural cell adhesion molecule 1 | ↓ | 0.10 | 4.08E-02 | | ↓ | 0.04 | 3.09E-02 |
| A0A0G2JSK1 | Serine proteinase inhibitor | ↓ | 0.10 | 4.10E-02 | | ↓ | 0.004 | 2.53E-02 |
| A0A0H2UHF8 | Orosomucoid 1 | ↓ | 0.10 | 2.09E-02 | | ↓ | 0.04 | 1.49E-02 |
| P07756 | Carbamoyl-phosphate synthase | ↓ | 0.11 | 3.15E-02 | | ↓ | 0.001 | 1.72E-02 |
| D3ZPV8 | Gamma-glutamyl cyclotransferase | ↓ | 0.11 | 2.43E-02 | | ↓ | 0.01 | 1.28E-02 |
| P05544 | Serine protease inhibitor A3L | ↓ | 0.11 | 1.60E-02 | | ↓ | 0.02 | 8.60E-03 |
| A0A0G2K135 | Complement factor I | ↓ | 0.11 | 4.52E-02 | | ↓ | 0.08 | 3.96E-02 |
| P28826 | Meprin A subunit beta | ↓ | 0.11 | 2.89E-02 | | ↓ | 0.07 | 2.23E-02 |
| Q5I0M1 | Beta-2-glycoprotein 1 | ↓ | 0.11 | 2.39E-02 | | ↓ | 0.03 | 1.35E-02 |
| P05545 | Serine protease inhibitor A3K | ↓ | 0.12 | 1.35E-02 | | ↓ | 0.04 | 7.61E-03 |
| P60711 | Actin, cytoplasmic 1 | ↓ | 0.12 | 2.96E-02 | | ↓ | 0.07 | 2.32E-02 |
| P52759 | 2-iminobutanoate/2-iminopropanoate deaminase | ↓ | 0.13 | 2.39E-02 | | ↓ | 0.01 | 1.02E-02 |
| P07154 | Procathepsin L | ↓ | 0.13 | 1.37E-02 | | ↓ | 0.04 | 6.77E-03 |
| D3ZAE6 | Vasorin | ↓ | 0.13 | 2.38E-02 | | ↓ | 0.05 | 1.41E-02 |
| Q8CG08 | Collagen triple helix repeat-containing protein 1 | ↓ | 0.13 | 2.81E-02 | | ↓ | 0.02 | 1.49E-02 |
| Q9QZH0 | Frizzled-4 | ↓ | 0.14 | 4.24E-02 | | ↓ | 0.08 | 3.30E-02 |
| P47967 | Galectin-5 | ↓ | 0.14 | 2.77E-02 | | ↓ | 0.05 | 1.65E-02 |
| Q01460 | Di-N-acetylchitobiase | ↓ | 0.15 | 4.97E-02 | | ↓ | 0.04 | 2.67E-02 |
| F1LR92 | Serpin family A member 3M | ↓ | 0.15 | 2.04E-02 | | ↓ | 0.04 | 8.53E-03 |
| A0A096MJP5 | Ectodysplasin A2 receptor | ↓ | 0.17 | 4.48E-02 | | ↓ | 0.04 | 2.20E-02 |
| A0A0G2K9B1 | Serine protease inhibitor | ↓ | 0.17 | 2.77E-02 | | ↓ | 0.04 | 1.05E-02 |
| F1MAE5 | Proline rich protein 2-like 1 | ↓ | 0.18 | 3.03E-02 | | ↓ | 0.14 | 2.47E-02 |
| G3V9L4 | Glycosylation-dependent cell adhesion molecule 1 | ↓ | 0.18 | 4.05E-02 | | ↓ | 0.01 | 1.25E-02 |
| Q32KJ5 | N-acetylglucosamine-6-sulfatase | ↓ | 0.18 | 1.91E-02 | | ↓ | 0.04 | 6.62E-03 |
| P08932 | T-kininogen 2 | ↓ | 0.19 | 4.31E-02 | | ↓ | 0.03 | 1.47E-02 |
| P24268 | Cathepsin D | ↓ | 0.20 | 3.40E-02 | | ↓ | 0.08 | 1.58E-02 |
| P01048 | T-kininogen 1 | ↓ | 0.22 | 2.67E-02 | | ↓ | 0.02 | 4.96E-03 |
| P98158 | Low-density lipoprotein receptor-related protein 2 | ↓ | 0.23 | 1.83E-02 | | ↓ | 0.13 | 8.81E-03 |
| D4A9V5 | Lysyl oxidase homolog | ↓ | 0.27 | 4.37E-02 | | ↓ | 0.04 | 6.31E-03 |
| P15684 | Aminopeptidase N | ↓ | 0.28 | 4.24E-02 | | ↓ | 0.10 | 1.32E-02 |
| A0A0G2K151 | Apolipoprotein E | ↓ | 0.29 | 4.66E-02 | | ↓ | 0.16 | 2.14E-02 |
| P08649 | Complement C4 | ↓ | 0.36 | 3.43E-02 | | ↓ | 0.15 | 3.65E-03 |
| Q6IRS6 | Fetuin B | ↓ | 0.40 | 1.27E-02 | | ↓ | 0.26 | 1.92E-03 |
| F1M698 | CD209f antigen | - | - | - | | ↓ | 0.001 | 3.38E-02 |
| P35213 | 14-3-3 protein beta/alpha | - | - | - | | ↓ | 0.0004 | 4.91E-02 |
| Q70Q42 | Resistin | - | - | - | | ↓ | 0.001 | 4.33E-02 |
| A0A0G2JU92 | Embigin | - | - | - | | ↓ | 0.001 | 4.21E-02 |
| P00762 | Serine protease 1 | - | - | - | | ↓ | 0.004 | 3.19E-02 |
| P0DP29 | Calmodulin-1 | - | - | - | | ↓ | 0.01 | 4.42E-02 |
| Q05175 | Brain acid soluble protein 1 | - | - | - | | ↓ | 0.01 | 4.08E-02 |
| D3ZIF6 | Urinary protein 1-like | - | - | - | | ↓ | 0.01 | 2.61E-02 |
| P10247 | H-2 class II histocompatibility antigen gamma chain | - | - | - | | ↓ | 0.01 | 3.36E-02 |
| P02625 | Parvalbumin alpha | - | - | - | | ↓ | 0.01 | 4.96E-02 |
| Q80YV8 | Stromal cell-derived factor 1 | - | - | - | | ↓ | 0.01 | 4.13E-02 |
| F1LML2 | Ubiquitin C | - | - | - | | ↓ | 0.01 | 3.95E-02 |
| D3ZJA4 | Seminal vesicle secretory protein 6 | - | - | - | | ↓ | 0.02 | 3.98E-02 |
| Q8CHN3 | WAP four-disulfide core domain protein 2 | - | - | - | | ↓ | 0.02 | 4.12E-02 |
| Q6AYS0 | Secreted and transmembrane 1A | - | - | - | | ↓ | 0.02 | 3.21E-02 |
| A0A0G2JWX9 | Chloride channel accessory 1 | - | - | - | | ↓ | 0.02 | 3.11E-02 |
| Q8R490 | Cadherin 13 | - | - | - | | ↓ | 0.02 | 3.27E-02 |
| D3Z841 | Butyrophilin like 10 | - | - | - | | ↓ | 0.02 | 6.89E-03 |
| M0R8H7 | Interferon alpha and beta receptor subunit 2 | - | - | - | | ↓ | 0.02 | 3.24E-02 |
| P97574 | Stanniocalcin-1 | - | - | - | | ↓ | 0.02 | 4.51E-02 |
| F1LPG1 | PBPC1BS-like | - | - | - | | ↓ | 0.02 | 3.40E-02 |
| D3ZJV7 | Lipocalin 4 | - | - | - | | ↓ | 0.02 | 2.57E-02 |
| A0A0G2JSQ7 | Kallikrein 1-related peptidase B3 | - | - | - | | ↓ | 0.02 | 3.14E-02 |
| F1LTN6 | Ig-like domain-containing protein | - | - | - | | ↓ | 0.02 | 1.19E-02 |
| A0A0G2JSJ7 | Carboxypeptidase Z | - | - | - | | ↓ | 0.02 | 2.28E-02 |
| P31044 | Phosphatidylethanolamine-binding protein 1 | - | - | - | | ↓ | 0.02 | 3.91E-02 |
| F1M9B2 | Insulin-like growth factor binding protein 7 | - | - | - | | ↓ | 0.03 | 4.46E-02 |
| E9PSP1 | Phospholipid transfer protein | - | - | - | | ↓ | 0.03 | 4.31E-02 |
| G3V9R2 | Complement factor H | - | - | - | | ↓ | 0.03 | 4.65E-02 |
| P27274 | CD59 glycoprotein | - | - | - | | ↓ | 0.03 | 2.17E-02 |
| F1LSA1 | CD44 antigen | - | - | - | | ↓ | 0.03 | 1.59E-02 |
| D3ZFP5 | Similar to Robo-1 | - | - | - | | ↓ | 0.03 | 1.97E-02 |
| P14841 | Cystatin-C | - | - | - | | ↓ | 0.03 | 3.63E-02 |
| A0A096P6L8 | Fibronectin 1 | - | - | - | | ↓ | 0.03 | 4.00E-02 |
| M0R7K9 | Common salivary protein 1 | - | - | - | | ↓ | 0.03 | 2.46E-02 |
| P53369 | Oxidized purine nucleoside triphosphate hydrolase | - | - | - | | ↓ | 0.03 | 4.69E-02 |
| Q499T2 | Gamma-interferon-inducible lysosomal thiol reductase | - | - | - | | ↓ | 0.04 | 4.73E-02 |
| A0A0G2JSG6 | Adenylate kinase 2 | - | - | - | | ↓ | 0.04 | 2.31E-02 |
| P48199 | C-reactive protein | - | - | - | | ↓ | 0.04 | 1.68E-03 |
| P81828 | Urinary protein 2 | - | - | - | | ↓ | 0.04 | 2.99E-02 |
| A0A0G2K1L0 | Tenascin C | - | - | - | | ↓ | 0.04 | 4.80E-02 |
| Q9WUC4 | Copper transport protein ATOX1 | - | - | - | | ↓ | 0.04 | 8.10E-03 |
| G3V7P2 | Fibrinogen-like 2 | - | - | - | | ↓ | 0.04 | 3.87E-02 |
| G3V615 | C3/C5 convertase | - | - | - | | ↓ | 0.04 | 2.74E-02 |
| P02761 | Major urinary protein | - | - | - | | ↓ | 0.04 | 1.69E-02 |
| Q64240 | Protein AMBP [Cleaved into: Alpha-1-microglobulin | - | - | - | | ↓ | 0.04 | 4.59E-02 |
| G3V6A0 | Platelet-derived growth factor receptor alpha | - | - | - | | ↓ | 0.04 | 1.43E-02 |
| A0A0G2JXI1 | Ac2-120 | - | - | - | | ↓ | 0.05 | 3.28E-02 |
| P02770 | Albumin | - | - | - | | ↓ | 0.05 | 1.66E-02 |
| M0R9L2 | Ciliary neurotrophic factor receptor | - | - | - | | ↓ | 0.05 | 2.44E-02 |
| Q9JHY1 | Junctional adhesion molecule A | - | - | - | | ↓ | 0.05 | 2.94E-02 |
| F1LPR6 | Immunoglobulin heavy constant epsilon | - | - | - | | ↓ | 0.05 | 4.27E-02 |
| P31211 | Corticosteroid-binding globulin | - | - | - | | ↓ | 0.05 | 3.55E-02 |
| Q9ESS6 | Basal cell adhesion molecule | - | - | - | | ↓ | 0.06 | 4.21E-02 |
| G3V6K1 | Transcobalamin 2 | - | - | - | | ↓ | 0.06 | 4.48E-02 |
| A0A0H2UHM3 | Haptoglobin | - | - | - | | ↓ | 0.06 | 4.90E-02 |
| P02781 | Prostatic steroid-binding protein C2 | - | - | - | | ↓ | 0.06 | 4.78E-02 |
| Q6DGG1 | Putative protein-lysine deacylase ABHD14B | - | - | - | | ↓ | 0.06 | 1.79E-02 |
| D3ZUR5 | Secreted LY6/PLAUR domain containing 2 | - | - | - | | ↓ | 0.07 | 4.24E-02 |
| P42854 | Regenerating islet-derived protein 3-gamma | - | - | - | | ↓ | 0.07 | 2.76E-02 |
| Q9JIK1 | Cadherin-related family member 5 | - | - | - | | ↓ | 0.07 | 4.84E-02 |
| Q5M7T5 | Antithrombin-III | - | - | - | | ↓ | 0.07 | 8.03E-03 |
| Q63041 | Alpha-1-macroglobulin | - | - | - | | ↓ | 0.07 | 8.50E-03 |
| Q9WUK5 | Inhibin beta C chain | - | - | - | | ↓ | 0.08 | 3.85E-02 |
| P21704 | Deoxyribonuclease-1 | - | - | - | | ↓ | 0.08 | 4.00E-02 |
| A0A0G2JVN2 | Secreted frizzled-related protein 4 | - | - | - | | ↓ | 0.08 | 4.77E-02 |
| A0A0G2K9X1 | Secreted phosphoprotein 24 | - | - | - | | ↓ | 0.09 | 1.88E-02 |
| Q9QX71 | Napsin | - | - | - | | ↓ | 0.09 | 1.59E-02 |
| Q6IRK9 | Carboxypeptidase Q | - | - | - | | ↓ | 0.09 | 2.00E-02 |
| A0A096MIX6 | Major urinary protein-like | - | - | - | | ↓ | 0.09 | 3.44E-02 |
| Q64230 | Meprin A subunit alpha | - | - | - | | ↓ | 0.09 | 2.00E-02 |
| A0A1W2Q642 | Peptidase inhibitor 16 | - | - | - | | ↓ | 0.10 | 2.28E-02 |
| P07171 | Calbindin | - | - | - | | ↓ | 0.10 | 4.41E-02 |
| F1LM84 | Nidogen 1 | - | - | - | | ↓ | 0.10 | 4.03E-02 |
| M0R7L1 | Lipase | - | - | - | | ↓ | 0.10 | 2.71E-02 |
| Q63257 | Interleukin-4 receptor subunit alpha | - | - | - | | ↓ | 0.11 | 2.10E-02 |
| A0A0G2JTX5 | Dipeptidyl peptidase 4 | - | - | - | | ↓ | 0.11 | 1.51E-02 |
| A0A0G2K890 | Ezrin | - | - | - | | ↓ | 0.12 | 3.23E-02 |
| A0A0G2JTC1 | Leukocyte immunoglobulin like receptor A5 | - | - | - | | ↓ | 0.12 | 4.49E-02 |
| D4ACM8 | Frizzled class receptor 7 | - | - | - | | ↓ | 0.12 | 1.98E-02 |
| F1LTJ5 | Heparan sulfate proteoglycan 2 | - | - | - | | ↓ | 0.13 | 1.81E-02 |
| A0A0G2K7Y9 | Zymogen granule protein 16B | - | - | - | | ↓ | 0.13 | 2.97E-02 |
| D3ZPC4 | L1 cell adhesion molecule | - | - | - | | ↓ | 0.14 | 4.51E-02 |
| G3V8Z5 | Ig-like domain-containing protein | - | - | - | | ↓ | 0.14 | 2.84E-02 |
| Q9EPB1 | Dipeptidyl peptidase 2 | - | - | - | | ↓ | 0.14 | 2.11E-02 |
| Q5EBA7 | Hepatocyte growth factor activator | - | - | - | | ↓ | 0.14 | 3.81E-02 |
| B2B9A9 | Ephrin B2 | - | - | - | | ↓ | 0.16 | 4.36E-02 |
| P04904 | Glutathione S-transferase alpha-3 | - | - | - | | ↓ | 0.17 | 2.96E-02 |
| G3V9R9 | Afamin | - | - | - | | ↓ | 0.18 | 2.92E-02 |
| D4A6I7 | Prostate stem cell antigen | - | - | - | | ↓ | 0.19 | 2.25E-02 |
| Q6AYS3 | Carboxypeptidase | - | - | - | | ↓ | 0.33 | 4.15E-02 |
| P02454 | Collagen alpha-1 | - | - | - | | ↓ | 0.40 | 4.34E-03 |

Table S2 Differential proteins produced by Day0 vs Day3 growth and development (FC≥1.5 or ≤0.67, P<0.05)

| Accession | Protein names | Trend | Fold change | P value |
| --- | --- | --- | --- | --- |
| G3V6P7 | Myosin, heavy chain 9 | ↓ | 0.4 | 1.14E-03 |
| B3EY86 | Lipocalin 11 | ↓ | 0.47 | 1.48E-02 |
| Q8K4G9 | Podocin | ↓ | 0.5 | 1.23E-02 |
| P84039 | Ectonucleotide pyrophosphatase/phosphodiesterase family member 5 | ↓ | 0.54 | 8.15E-04 |
| A0A140TAG4 | NPHS1 adhesion molecule, nephrin | ↓ | 0.61 | 2.41E-02 |
| Q642A7 | Protein FAM151A | ↓ | 0.63 | 8.56E-04 |
| F1MA37 | Serine protease 8 | ↓ | 0.65 | 1.02E-03 |
| O54861 | Sortilin | ↓ | 0.66 | 8.86E-03 |
| M0R5R0 | Protein S | ↑ | 1.5 | 3.03E-02 |
| P24268 | Cathepsin D | ↑ | 1.54 | 3.05E-03 |
| Q498R7 | CXXC motif containing zinc binding protein | ↑ | 1.54 | 4.07E-02 |
| A0A0G2QC04 | Plastin 1 | ↑ | 1.57 | 1.17E-02 |
| P18418 | Calreticulin | ↑ | 1.6 | 4.26E-02 |
| O88989 | Malate dehydrogenase, cytoplasmic | ↑ | 1.63 | 2.46E-02 |
| P07379 | Phosphoenolpyruvate carboxykinase, cytosolic | ↑ | 1.63 | 2.84E-02 |
| Q5U355 | Integrin alpha FG-GAP repeat containing 1 | ↑ | 1.63 | 4.19E-02 |
| A0A0G2K8I5 | Protocadherin 19 | ↑ | 1.64 | 1.61E-02 |
| A0A0G2JTX5 | Dipeptidyl peptidase 4 | ↑ | 1.67 | 2.17E-02 |
| P45592 | Cofilin-1 | ↑ | 1.68 | 2.39E-02 |
| A0A0G2JSS8 | Peroxiredoxin-5 | ↑ | 1.68 | 3.86E-02 |
| A0A0G2JV31 | X-prolyl aminopeptidase | ↑ | 1.69 | 1.10E-02 |
| A0A0G2K6T9 | Protocadherin 1 | ↑ | 1.71 | 2.37E-02 |
| D3ZUD8 | Transmembrane 9 superfamily member | ↑ | 1.73 | 2.51E-02 |
| A0A0G2K6T8 | - | ↑ | 1.9 | 3.19E-02 |
| Q6IRE4 | Tumor susceptibility gene 101 protein | ↑ | 1.95 | 1.73E-02 |
| Q9WUK5 | Inhibin beta C chain | ↑ | 2.01 | 3.26E-02 |
| P24368 | Peptidyl-prolyl cis-trans isomerase B | ↑ | 2.04 | 2.56E-02 |
| Q6P6V0 | Glucose-6-phosphate isomerase | ↑ | 2.15 | 1.87E-02 |
| P63322 | Ras-related protein Ral-A | ↑ | 2.19 | 4.06E-02 |
| Q9R0T3 | DnaJ homolog subfamily C member 3 | ↑ | 2.23 | 3.49E-02 |
| G3V8J3 | Chymotrypsin-like | ↑ | 2.45 | 1.47E-02 |
| A0A0G2JXJ3 | FAM3 metabolism regulating signaling molecule D | ↑ | 2.79 | 2.88E-02 |
| G3V8K5 | Growth differentiation Factor 15 | ↑ | 2.86 | 3.91E-02 |
| F1M498 | Gastrokine 3 | ↑ | 2.94 | 3.28E-02 |
| Q9QZK9 | Deoxyribonuclease-2-beta | ↑ | 3.25 | 1.91E-02 |
| Q80YV8 | Stromal cell-derived Factor 1 | ↑ | 3.27 | 4.66E-02 |
| A0A0G2K5X1 | Putative lysozyme C-2 | ↑ | 3.53 | 2.25E-02 |

Table S3 Day3 vs Day6 Differential Proteins Produced by Growth and Development (FC≥1.5 or ≤0.67, P<0.05)

| Accession | Protein names | Trend | Fold change | P value |
| --- | --- | --- | --- | --- |
| Q5I0D7 | Xaa-Pro dipeptidase | ↓ | 0.23 | 4.41E-02 |
| Q812E9 | Neuronal membrane glycoprotein M6-a | ↓ | 0.31 | 8.50E-04 |
| Q6MG71 | Choline transporter-like protein 4 | ↓ | 0.32 | 2.48E-02 |
| B0BNJ1 | LOC683667 protein | ↓ | 0.36 | 3.45E-02 |
| G3V8L3 | Lamin A, isoform CRA_b | ↓ | 0.38 | 4.64E-02 |
| G3V847 | Sodium-dependent neutral amino acid transporter B(0)AT3 | ↓ | 0.39 | 1.22E-03 |
| D3ZLA3 | Copine 3 | ↓ | 0.39 | 6.11E-03 |
| P53790 | Sodium/glucose cotransporter 1 | ↓ | 0.39 | 1.37E-02 |
| D4A8C5 | 1-phosphatidylinositol 4,5-bisphosphate phosphodiesterase | ↓ | 0.39 | 4.40E-02 |
| Q9JJ19 | Na(+)/H(+) exchange regulatory cofactor NHE-RF1 | ↓ | 0.4 | 2.42E-03 |
| F1LVR0 | IgLON family member 5 | ↓ | 0.4 | 3.13E-02 |
| Q80W57 | Broad substrate specificity ATP-binding cassette transporter ABCG2 | ↓ | 0.41 | 2.53E-03 |
| A0A0G2K595 | Solute carrier family 23 member 1 | ↓ | 0.42 | 5.28E-03 |
| P51607 | N-acylglucosamine 2-epimerase | ↓ | 0.42 | 3.76E-02 |
| P07314 | Glutathione hydrolase 1 proenzyme | ↓ | 0.43 | 6.42E-05 |
| Q5FVI6 | V-type proton ATPase subunit C 1 | ↓ | 0.44 | 4.10E-02 |
| A0A0A0MXX5 | 3-hydroxybutyrate dehydrogenase 2 | ↓ | 0.45 | 1.60E-02 |
| D3ZV91 | trans-L-3-hydroxyproline dehydratase | ↓ | 0.45 | 3.61E-02 |
| Q9JJ40 | Na(+)/H(+) exchange regulatory cofactor NHE-RF3 | ↓ | 0.46 | 5.04E-03 |
| Q499Q4 | Phosphoglucomutase 1 | ↓ | 0.46 | 4.37E-02 |
| G3V8X5 | Solute carrier family 5 | ↓ | 0.47 | 1.39E-02 |
| A0A0G2JT43 | Solute carrier family 2, facilitated glucose transporter member 5 | ↓ | 0.47 | 1.74E-02 |
| P09606 | Glutamine synthetase | ↓ | 0.47 | 2.69E-02 |
| E9PU23 | Delta-like protein | ↓ | 0.47 | 3.45E-02 |
| A0A0G2JWD0 | Prominin 1 | ↓ | 0.48 | 1.47E-02 |
| A1L1J8 | RAB5B, member RAS oncogene family | ↓ | 0.48 | 3.64E-02 |
| Q641Z6 | EH domain-containing protein 1 | ↓ | 0.49 | 2.75E-03 |
| G3V6A0 | Platelet-derived growth factor receptor alpha | ↓ | 0.5 | 1.18E-02 |
| P48508 | Glutamate--cysteine ligase regulatory subunit | ↓ | 0.5 | 1.25E-02 |
| Q9WUW9 | Sulfotransferase 1C2A | ↓ | 0.5 | 2.55E-02 |
| A0A0G2JSV5 | 3-hydroxyanthranilate 3,4-dioxygenase | ↓ | 0.51 | 2.47E-02 |
| G3V6D9 | Na(+)/H(+) exchange regulatory cofactor NHE-RF | ↓ | 0.51 | 3.61E-02 |
| P16975 | SPARC | ↓ | 0.51 | 4.72E-02 |
| Q5M876 | N-acyl-aromatic-L-amino acid amidohydrolase | ↓ | 0.52 | 4.71E-03 |
| A0A0A0MXT4 | Solute carrier organic anion transporter family member | ↓ | 0.53 | 1.95E-03 |
| P61459 | Pterin-4-alpha-carbinolamine dehydratase | ↓ | 0.53 | 1.15E-02 |
| P02770 | Albumin | ↓ | 0.53 | 3.80E-02 |
| D3ZXY4 | Aldehyde dehydrogenase 8 family, member A1 | ↓ | 0.53 | 3.81E-02 |
| P80254 | D-dopachrome decarboxylase | ↓ | 0.55 | 1.55E-02 |
| Q64602 | Kynurenine/alpha-aminoadipate aminotransferase, mitochondrial | ↓ | 0.55 | 3.66E-02 |
| A0A0G2JSQ7 | Kallikrein 1-related peptidase B3 | ↓ | 0.55 | 3.86E-02 |
| P97546 | Neuroplastin | ↓ | 0.57 | 2.39E-02 |
| P32755 | 4-hydroxyphenylpyruvate dioxygenase | ↓ | 0.57 | 4.85E-02 |
| Q3KRC4 | G-protein coupled receptor family C Group 5 member C | ↓ | 0.6 | 4.10E-02 |
| A0A0G2QC04 | Plastin 1 | ↓ | 0.61 | 7.25E-03 |
| G3V7L4 | Cadherin 16 | ↓ | 0.61 | 1.86E-02 |
| A0A0G2JSJ2 | Cytidine/uridine monophosphate kinase 1 | ↓ | 0.62 | 2.12E-03 |
| F1LQI1 | Hydroxyacyl glutathione hydrolase | ↓ | 0.62 | 2.03E-02 |
| Q5U362 | Annexin | ↓ | 0.62 | 2.43E-02 |
| Q4V7D9 | Acid sphingomyelinase-like phosphodiesterase | ↓ | 0.62 | 4.24E-02 |
| Q5M8C3 | Serine | ↓ | 0.63 | 1.04E-02 |
| P38918 | Aflatoxin B1 aldehyde reductase member 3 | ↓ | 0.63 | 3.93E-02 |
| F1MAD3 | Polycystin 1, transient receptor potential channel interacting | ↓ | 0.64 | 1.13E-02 |
| P04904 | Glutathione S-transferase alpha-3 | ↓ | 0.65 | 2.75E-02 |
| Q6AXM6 | Intercellular adhesion molecule 2 | ↓ | 0.65 | 3.84E-02 |
| D3ZE16 | Alpha-L-iduronidase | ↓ | 0.66 | 4.13E-03 |
| A0A140TAG4 | NPHS1 adhesion molecule, nephrin | ↓ | 0.66 | 7.13E-03 |
| Q6Q0N1 | Cytosolic nonspecific dipeptidase | ↓ | 0.66 | 1.23E-02 |
| F1M978 | Inositol-1-monophosphatase | ↓ | 0.66 | 1.69E-02 |
| Q05030 | Platelet-derived growth factor receptor beta | ↓ | 0.66 | 4.40E-02 |
| A0A0G2KB26 | Matrix remodeling-associated protein 8 | ↑ | 1.53 | 2.40E-03 |
| D3ZH39 | receptor protein-tyrosine kinase | ↑ | 1.57 | 9.17E-03 |
| A0A0G2JZ18 | Pentraxin 4 | ↑ | 1.58 | 1.11E-02 |
| D4AA31 | Prolylcarboxypeptidase | ↑ | 1.66 | 3.81E-03 |
| D4A4U3 | Magnesium-dependent phosphatase 1 | ↑ | 1.68 | 2.92E-02 |
| G3V7D0 | Matrix metallopeptidase 8 | ↑ | 1.69 | 6.91E-03 |
| Q63041 | Alpha-1-macroglobulin | ↑ | 1.71 | 2.89E-04 |
| P50137 | Transketolase | ↑ | 1.76 | 2.26E-02 |
| F7F389 | Complement C9 | ↑ | 1.82 | 4.22E-03 |
| P01681 | Ig kappa chain V region S211 | ↑ | 1.93 | 3.36E-02 |
| D4A400 | Lactoperoxidase | ↑ | 1.96 | 9.70E-03 |
| Q5I0E1 | Leucine-rich alpha-2-glycoprotein 1 | ↑ | 2 | 4.74E-03 |
| D3ZUQ1 | Lipase | ↑ | 2.05 | 8.91E-04 |
| M0R7L1 | Lipase | ↑ | 2.18 | 1.22E-03 |
| P14668 | Annexin A5 | ↑ | 2.3 | 1.21E-03 |

Table S4 Biological Processes Enriched for Differential Proteins Produced under Relaxed Conditions Compared to Day0 vs. Day3 (P<0.05)

| Biological process | P-Value |
| --- | --- |
| cellular response to cytochalasin B | 2.20E-10 |
| negative regulation of endopeptidase activity | 6.90E-10 |
| regulation of norepinephrine uptake | 1.50E-09 |
| protein localization to adherens junction | 5.40E-09 |
| morphogenesis of a polarized epithelium | 1.40E-08 |
| regulation of transmembrane transporter activity | 3.10E-08 |
| adherens junction assembly | 5.80E-08 |
| apical protein localization | 1.60E-07 |
| histone H2A acetylation | 2.00E-07 |
| cellular response to electrical stimulus | 3.70E-07 |
| inflammatory response | 4.50E-07 |
| postsynaptic actin cytoskeleton organization | 1.10E-06 |
| regulation of protein localization to plasma membrane | 2.10E-06 |
| regulation of cyclin-dependent protein serine/threonine kinase activity | 3.30E-06 |
| cellular phosphate ion homeostasis | 3.80E-06 |
| histone H4 acetylation | 4.50E-06 |
| positive regulation of double-strand break repair via homologous recombination | 5.50E-06 |
| cell motility | 9.20E-06 |
| response to immobilization stress | 1.30E-05 |
| establishment or maintenance of cell polarity | 1.30E-05 |
| synaptic vesicle endocytosis | 1.60E-05 |
| aging | 3.50E-05 |
| negative regulation of protein binding | 6.90E-05 |
| collecting duct development | 1.00E-04 |
| retina homeostasis | 1.30E-04 |
| acute-phase response | 2.10E-04 |
| retina development in camera-type eye | 2.90E-04 |
| axonogenesis | 3.10E-04 |
| response to mechanical stimulus | 3.50E-04 |
| negative regulation of blood coagulation | 5.30E-04 |
| circadian rhythm | 5.30E-04 |
| vasodilation | 7.10E-04 |
| cellular chloride ion homeostasis | 8.10E-04 |
| response to lipopolysaccharide | 1.00E-03 |
| positive regulation of MAPK cascade | 2.30E-03 |
| protein processing | 2.30E-03 |
| cellular sodium ion homeostasis | 2.70E-03 |
| osteoblast differentiation | 3.10E-03 |
| proteolysis | 3.10E-03 |
| cell adhesion | 3.20E-03 |
| response to xenobiotic stimulus | 3.50E-03 |
| response to vitamin D | 3.70E-03 |
| cellular calcium ion homeostasis | 3.80E-03 |
| negative regulation of collateral sprouting of intact axon in response to injury | 5.30E-03 |
| negative regulation of canonical Wnt signaling pathway | 5.90E-03 |
| regulation of cell cycle | 7.50E-03 |
| Factor XII activation | 1.10E-02 |
| response to steroid hormone | 1.20E-02 |
| folic acid import into cell | 1.30E-02 |
| androgen catabolic process | 1.30E-02 |
| response to organic substance | 1.40E-02 |
| lipoprotein catabolic process | 1.60E-02 |
| kidney development | 1.60E-02 |
| neutrophil chemotaxis | 1.70E-02 |
| positive regulation of estradiol secretion | 1.80E-02 |
| urate biosynthetic process | 1.80E-02 |
| negative regulation of cholesterol biosynthetic process | 2.10E-02 |
| response to caloric restriction | 2.10E-02 |
| regulation of blood pressure | 2.10E-02 |
| protein import | 2.40E-02 |
| gene expression | 2.40E-02 |
| metanephric proximal tubule development | 2.60E-02 |
| negative regulation of lymphocyte proliferation | 2.60E-02 |
| animal organ regeneration | 2.70E-02 |
| sensory perception of sound | 2.70E-02 |
| homotypic cell-cell adhesion | 3.10E-02 |
| peripheral nervous system axon regeneration | 3.10E-02 |
| protein catabolic process | 3.30E-02 |
| beta-amyloid clearance | 3.60E-02 |
| cellular response to interleukin-1 | 3.90E-02 |
| cellular response to interferon-gamma | 4.60E-02 |
| hepatocyte differentiation | 4.90E-02 |
| multicellular organismal response to stress | 4.90E-02 |

Table S5 Signaling Pathways Enriched for Differential Proteins Produced under Relaxed Conditions Compared to Day0 vs. Day3 (P<0.05)

| pathway | [P-Value](https://david.ncifcrf.gov/chartReport.jsp?d-16544-p=1&d-16544-o=1&annot=52&d-16544-s=7" \o "https://david.ncifcrf.gov/chartReport.jsp?d-16544-p=1&d-16544-o=1&annot=52&d-16544-s=7) |
| --- | --- |
| Apoptosis | 1.10E-04 |
| Phagosome | 4.60E-04 |
| Proteoglycans in cancer | 7.50E-04 |
| Leukocyte transendothelial migration | 7.90E-04 |
| Regulation of actin cytoskeleton | 1.30E-03 |
| Fluid shear stress and atherosclerosis | 1.90E-03 |
| Hippo signaling pathway | 2.20E-03 |
| Gastric acid secretion | 2.40E-03 |
| Bacterial invasion of epithelial cells | 2.60E-03 |
| Arrhythmogenic right ventricular cardiomyopathy | 2.60E-03 |
| Tight junction | 2.90E-03 |
| Viral myocarditis | 3.40E-03 |
| Complement and coagulation cascades | 3.50E-03 |
| Hepatocellular carcinoma | 3.60E-03 |
| Hypertrophic cardiomyopathy | 4.10E-03 |
| Adherens junction | 4.40E-03 |
| Dilated cardiomyopathy | 4.50E-03 |
| Focal adhesion | 5.40E-03 |
| ATP-dependent chromatin remodeling | 9.30E-03 |
| Thyroid hormone signaling pathway | 9.70E-03 |
| Platelet activation | 9.90E-03 |
| Sphingolipid signaling pathway | 9.90E-03 |
| Yersinia infection | 1.20E-02 |
| Oxytocin signaling pathway | 1.70E-02 |
| Influenza A | 2.20E-02 |
| Cell adhesion molecules | 2.60E-02 |
| Glutathione metabolism | 2.70E-02 |
| Neutrophil extracellular trap formation | 2.80E-02 |
| Motor proteins | 3.10E-02 |
| Rap1 signaling pathway | 4.00E-02 |
| Amyotrophic lateral sclerosis | 4.10E-02 |

Table S6 Biological Processes Produced by Differential Protein Enrichment to Day0 vs. Day4 Compared to Relaxed Conditions (P<0.05)

| Biological process | [P-Value](https://david.ncifcrf.gov/chartReport.jsp?d-16544-p=1&d-16544-o=1&annot=27&d-16544-s=7" \o "https://david.ncifcrf.gov/chartReport.jsp?d-16544-p=1&d-16544-o=1&annot=27&d-16544-s=7) |
| --- | --- |
| [negative regulation of endopeptidase activity](http://www.ebi.ac.uk/QuickGO/GTerm?id=GO:0010951" \o "http://www.ebi.ac.uk/QuickGO/GTerm?id=GO:0010951) | 1.20E-09 |
| [proteolysis](http://www.ebi.ac.uk/QuickGO/GTerm?id=GO:0006508" \o "http://www.ebi.ac.uk/QuickGO/GTerm?id=GO:0006508) | 5.50E-09 |
| [cellular response to cytochalasin B](http://www.ebi.ac.uk/QuickGO/GTerm?id=GO:0072749" \o "http://www.ebi.ac.uk/QuickGO/GTerm?id=GO:0072749) | 1.00E-08 |
| [aging](http://www.ebi.ac.uk/QuickGO/GTerm?id=GO:0007568" \o "http://www.ebi.ac.uk/QuickGO/GTerm?id=GO:0007568) | 1.50E-08 |
| [acute-phase response](http://www.ebi.ac.uk/QuickGO/GTerm?id=GO:0006953" \o "http://www.ebi.ac.uk/QuickGO/GTerm?id=GO:0006953) | 1.80E-08 |
| [regulation of norepinephrine uptake](http://www.ebi.ac.uk/QuickGO/GTerm?id=GO:0051621" \o "http://www.ebi.ac.uk/QuickGO/GTerm?id=GO:0051621) | 7.00E-08 |
| [protein localization to adherens junction](http://www.ebi.ac.uk/QuickGO/GTerm?id=GO:0071896" \o "http://www.ebi.ac.uk/QuickGO/GTerm?id=GO:0071896) | 2.50E-07 |
| [cell adhesion](http://www.ebi.ac.uk/QuickGO/GTerm?id=GO:0007155" \o "http://www.ebi.ac.uk/QuickGO/GTerm?id=GO:0007155) | 3.10E-07 |
| [morphogenesis of a polarized epithelium](http://www.ebi.ac.uk/QuickGO/GTerm?id=GO:0001738" \o "http://www.ebi.ac.uk/QuickGO/GTerm?id=GO:0001738) | 6.50E-07 |
| [regulation of transmembrane transporter activity](http://www.ebi.ac.uk/QuickGO/GTerm?id=GO:0022898" \o "http://www.ebi.ac.uk/QuickGO/GTerm?id=GO:0022898) | 1.40E-06 |
| [postsynaptic actin cytoskeleton organization](http://www.ebi.ac.uk/QuickGO/GTerm?id=GO:0098974" \o "http://www.ebi.ac.uk/QuickGO/GTerm?id=GO:0098974) | 1.70E-06 |
| [response to mechanical stimulus](http://www.ebi.ac.uk/QuickGO/GTerm?id=GO:0009612" \o "http://www.ebi.ac.uk/QuickGO/GTerm?id=GO:0009612) | 2.50E-06 |
| [adherens junction assembly](http://www.ebi.ac.uk/QuickGO/GTerm?id=GO:0034333" \o "http://www.ebi.ac.uk/QuickGO/GTerm?id=GO:0034333) | 2.60E-06 |
| [retina homeostasis](http://www.ebi.ac.uk/QuickGO/GTerm?id=GO:0001895" \o "http://www.ebi.ac.uk/QuickGO/GTerm?id=GO:0001895) | 4.90E-06 |
| [embryo implantation](http://www.ebi.ac.uk/QuickGO/GTerm?id=GO:0007566" \o "http://www.ebi.ac.uk/QuickGO/GTerm?id=GO:0007566) | 5.20E-06 |
| [response to organic substance](http://www.ebi.ac.uk/QuickGO/GTerm?id=GO:0010033" \o "http://www.ebi.ac.uk/QuickGO/GTerm?id=GO:0010033) | 5.60E-06 |
| [apical protein localization](http://www.ebi.ac.uk/QuickGO/GTerm?id=GO:0045176" \o "http://www.ebi.ac.uk/QuickGO/GTerm?id=GO:0045176) | 7.30E-06 |
| [inflammatory response](http://www.ebi.ac.uk/QuickGO/GTerm?id=GO:0006954" \o "http://www.ebi.ac.uk/QuickGO/GTerm?id=GO:0006954) | 7.90E-06 |
| [histone H2A acetylation](http://www.ebi.ac.uk/QuickGO/GTerm?id=GO:0043968" \o "http://www.ebi.ac.uk/QuickGO/GTerm?id=GO:0043968) | 9.10E-06 |
| [cellular response to electrical stimulus](http://www.ebi.ac.uk/QuickGO/GTerm?id=GO:0071257" \o "http://www.ebi.ac.uk/QuickGO/GTerm?id=GO:0071257) | 1.60E-05 |
| [response to xenobiotic stimulus](http://www.ebi.ac.uk/QuickGO/GTerm?id=GO:0009410" \o "http://www.ebi.ac.uk/QuickGO/GTerm?id=GO:0009410) | 2.90E-05 |
| [cytokine-mediated signaling pathway](http://www.ebi.ac.uk/QuickGO/GTerm?id=GO:0019221" \o "http://www.ebi.ac.uk/QuickGO/GTerm?id=GO:0019221) | 6.30E-05 |
| [cellular phosphate ion homeostasis](http://www.ebi.ac.uk/QuickGO/GTerm?id=GO:0030643" \o "http://www.ebi.ac.uk/QuickGO/GTerm?id=GO:0030643) | 6.50E-05 |
| [zymogen activation](http://www.ebi.ac.uk/QuickGO/GTerm?id=GO:0031638" \o "http://www.ebi.ac.uk/QuickGO/GTerm?id=GO:0031638) | 8.10E-05 |
| [regulation of protein localization to plasma membrane](http://www.ebi.ac.uk/QuickGO/GTerm?id=GO:1903076" \o "http://www.ebi.ac.uk/QuickGO/GTerm?id=GO:1903076) | 9.10E-05 |
| [regulation of cyclin-dependent protein serine/threonine kinase activity](http://www.ebi.ac.uk/QuickGO/GTerm?id=GO:0000079" \o "http://www.ebi.ac.uk/QuickGO/GTerm?id=GO:0000079) | 1.40E-04 |
| [retina development in camera-type eye](http://www.ebi.ac.uk/QuickGO/GTerm?id=GO:0060041" \o "http://www.ebi.ac.uk/QuickGO/GTerm?id=GO:0060041) | 1.80E-04 |
| [histone H4 acetylation](http://www.ebi.ac.uk/QuickGO/GTerm?id=GO:0043967" \o "http://www.ebi.ac.uk/QuickGO/GTerm?id=GO:0043967) | 1.90E-04 |
| [positive regulation of double-strand break repair via homologous recombination](http://www.ebi.ac.uk/QuickGO/GTerm?id=GO:1905168" \o "http://www.ebi.ac.uk/QuickGO/GTerm?id=GO:1905168) | 2.20E-04 |
| [immune response](http://www.ebi.ac.uk/QuickGO/GTerm?id=GO:0006955" \o "http://www.ebi.ac.uk/QuickGO/GTerm?id=GO:0006955) | 2.60E-04 |
| [response to nutrient levels](http://www.ebi.ac.uk/QuickGO/GTerm?id=GO:0031667" \o "http://www.ebi.ac.uk/QuickGO/GTerm?id=GO:0031667) | 3.50E-04 |
| [cell motility](http://www.ebi.ac.uk/QuickGO/GTerm?id=GO:0048870" \o "http://www.ebi.ac.uk/QuickGO/GTerm?id=GO:0048870) | 3.70E-04 |
| [response to organic cyclic compound](http://www.ebi.ac.uk/QuickGO/GTerm?id=GO:0014070" \o "http://www.ebi.ac.uk/QuickGO/GTerm?id=GO:0014070) | 4.70E-04 |
| [response to immobilization stress](http://www.ebi.ac.uk/QuickGO/GTerm?id=GO:0035902" \o "http://www.ebi.ac.uk/QuickGO/GTerm?id=GO:0035902) | 5.00E-04 |
| [animal organ regeneration](http://www.ebi.ac.uk/QuickGO/GTerm?id=GO:0031100" \o "http://www.ebi.ac.uk/QuickGO/GTerm?id=GO:0031100) | 5.30E-04 |
| [establishment or maintenance of cell polarity](http://www.ebi.ac.uk/QuickGO/GTerm?id=GO:0007163" \o "http://www.ebi.ac.uk/QuickGO/GTerm?id=GO:0007163) | 5.30E-04 |
| [complement activation](http://www.ebi.ac.uk/QuickGO/GTerm?id=GO:0006956" \o "http://www.ebi.ac.uk/QuickGO/GTerm?id=GO:0006956) | 5.60E-04 |
| [platelet aggregation](http://www.ebi.ac.uk/QuickGO/GTerm?id=GO:0070527" \o "http://www.ebi.ac.uk/QuickGO/GTerm?id=GO:0070527) | 5.70E-04 |
| [synaptic vesicle endocytosis](http://www.ebi.ac.uk/QuickGO/GTerm?id=GO:0048488" \o "http://www.ebi.ac.uk/QuickGO/GTerm?id=GO:0048488) | 6.10E-04 |
| [negative regulation of cell adhesion](http://www.ebi.ac.uk/QuickGO/GTerm?id=GO:0007162" \o "http://www.ebi.ac.uk/QuickGO/GTerm?id=GO:0007162) | 6.50E-04 |
| [collecting duct development](http://www.ebi.ac.uk/QuickGO/GTerm?id=GO:0072044" \o "http://www.ebi.ac.uk/QuickGO/GTerm?id=GO:0072044) | 6.80E-04 |
| [protein catabolic process](http://www.ebi.ac.uk/QuickGO/GTerm?id=GO:0030163" \o "http://www.ebi.ac.uk/QuickGO/GTerm?id=GO:0030163) | 8.20E-04 |
| [complement activation, classical pathway](http://www.ebi.ac.uk/QuickGO/GTerm?id=GO:0006958" \o "http://www.ebi.ac.uk/QuickGO/GTerm?id=GO:0006958) | 9.50E-04 |
| [vasodilation](http://www.ebi.ac.uk/QuickGO/GTerm?id=GO:0042311" \o "http://www.ebi.ac.uk/QuickGO/GTerm?id=GO:0042311) | 1.10E-03 |
| [positive regulation of calcium-mediated signaling](http://www.ebi.ac.uk/QuickGO/GTerm?id=GO:0050850" \o "http://www.ebi.ac.uk/QuickGO/GTerm?id=GO:0050850) | 1.10E-03 |
| [leukocyte cell-cell adhesion](http://www.ebi.ac.uk/QuickGO/GTerm?id=GO:0007159" \o "http://www.ebi.ac.uk/QuickGO/GTerm?id=GO:0007159) | 1.20E-03 |
| [response to hypoxia](http://www.ebi.ac.uk/QuickGO/GTerm?id=GO:0001666" \o "http://www.ebi.ac.uk/QuickGO/GTerm?id=GO:0001666) | 1.40E-03 |
| [response to vitamin D](http://www.ebi.ac.uk/QuickGO/GTerm?id=GO:0033280" \o "http://www.ebi.ac.uk/QuickGO/GTerm?id=GO:0033280) | 1.60E-03 |
| [integrin activation](http://www.ebi.ac.uk/QuickGO/GTerm?id=GO:0033622" \o "http://www.ebi.ac.uk/QuickGO/GTerm?id=GO:0033622) | 2.40E-03 |
| [negative regulation of protein binding](http://www.ebi.ac.uk/QuickGO/GTerm?id=GO:0032091" \o "http://www.ebi.ac.uk/QuickGO/GTerm?id=GO:0032091) | 2.50E-03 |
| [positive regulation of fibroblast proliferation](http://www.ebi.ac.uk/QuickGO/GTerm?id=GO:0048146" \o "http://www.ebi.ac.uk/QuickGO/GTerm?id=GO:0048146) | 2.80E-03 |
| [positive regulation of endothelial cell proliferation](http://www.ebi.ac.uk/QuickGO/GTerm?id=GO:0001938" \o "http://www.ebi.ac.uk/QuickGO/GTerm?id=GO:0001938) | 2.80E-03 |
| [peripheral nervous system axon regeneration](http://www.ebi.ac.uk/QuickGO/GTerm?id=GO:0014012" \o "http://www.ebi.ac.uk/QuickGO/GTerm?id=GO:0014012) | 2.90E-03 |
| [homotypic cell-cell adhesion](http://www.ebi.ac.uk/QuickGO/GTerm?id=GO:0034109" \o "http://www.ebi.ac.uk/QuickGO/GTerm?id=GO:0034109) | 2.90E-03 |
| [positive regulation of MAPK cascade](http://www.ebi.ac.uk/QuickGO/GTerm?id=GO:0043410" \o "http://www.ebi.ac.uk/QuickGO/GTerm?id=GO:0043410) | 3.10E-03 |
| [negative regulation of blood coagulation](http://www.ebi.ac.uk/QuickGO/GTerm?id=GO:0030195" \o "http://www.ebi.ac.uk/QuickGO/GTerm?id=GO:0030195) | 3.40E-03 |
| [positive regulation of cell migration](http://www.ebi.ac.uk/QuickGO/GTerm?id=GO:0030335" \o "http://www.ebi.ac.uk/QuickGO/GTerm?id=GO:0030335) | 3.60E-03 |
| [wound healing](http://www.ebi.ac.uk/QuickGO/GTerm?id=GO:0042060" \o "http://www.ebi.ac.uk/QuickGO/GTerm?id=GO:0042060) | 3.60E-03 |
| [response to lipopolysaccharide](http://www.ebi.ac.uk/QuickGO/GTerm?id=GO:0032496" \o "http://www.ebi.ac.uk/QuickGO/GTerm?id=GO:0032496) | 3.90E-03 |
| [response to heat](http://www.ebi.ac.uk/QuickGO/GTerm?id=GO:0009408" \o "http://www.ebi.ac.uk/QuickGO/GTerm?id=GO:0009408) | 4.00E-03 |
| [positive regulation of cell-substrate adhesion](http://www.ebi.ac.uk/QuickGO/GTerm?id=GO:0010811" \o "http://www.ebi.ac.uk/QuickGO/GTerm?id=GO:0010811) | 4.30E-03 |
| [cellular response to fibroblast growth factor stimulus](http://www.ebi.ac.uk/QuickGO/GTerm?id=GO:0044344" \o "http://www.ebi.ac.uk/QuickGO/GTerm?id=GO:0044344) | 4.30E-03 |
| [protein processing](http://www.ebi.ac.uk/QuickGO/GTerm?id=GO:0016485" \o "http://www.ebi.ac.uk/QuickGO/GTerm?id=GO:0016485) | 4.70E-03 |
| [cellular chloride ion homeostasis](http://www.ebi.ac.uk/QuickGO/GTerm?id=GO:0030644" \o "http://www.ebi.ac.uk/QuickGO/GTerm?id=GO:0030644) | 5.20E-03 |
| [osteoblast differentiation](http://www.ebi.ac.uk/QuickGO/GTerm?id=GO:0001649" \o "http://www.ebi.ac.uk/QuickGO/GTerm?id=GO:0001649) | 6.80E-03 |
| [response to thyroid hormone](http://www.ebi.ac.uk/QuickGO/GTerm?id=GO:0097066" \o "http://www.ebi.ac.uk/QuickGO/GTerm?id=GO:0097066) | 7.30E-03 |
| [negative regulation of cell-substrate adhesion](http://www.ebi.ac.uk/QuickGO/GTerm?id=GO:0010812" \o "http://www.ebi.ac.uk/QuickGO/GTerm?id=GO:0010812) | 7.30E-03 |
| [epithelial cell differentiation](http://www.ebi.ac.uk/QuickGO/GTerm?id=GO:0030855" \o "http://www.ebi.ac.uk/QuickGO/GTerm?id=GO:0030855) | 7.90E-03 |
| [cellular response to interleukin-1](http://www.ebi.ac.uk/QuickGO/GTerm?id=GO:0071347" \o "http://www.ebi.ac.uk/QuickGO/GTerm?id=GO:0071347) | 8.60E-03 |
| [cellular calcium ion homeostasis](http://www.ebi.ac.uk/QuickGO/GTerm?id=GO:0006874" \o "http://www.ebi.ac.uk/QuickGO/GTerm?id=GO:0006874) | 8.60E-03 |
| [response to steroid hormone](http://www.ebi.ac.uk/QuickGO/GTerm?id=GO:0048545" \o "http://www.ebi.ac.uk/QuickGO/GTerm?id=GO:0048545) | 9.50E-03 |
| [regulation of cell growth](http://www.ebi.ac.uk/QuickGO/GTerm?id=GO:0001558" \o "http://www.ebi.ac.uk/QuickGO/GTerm?id=GO:0001558) | 9.50E-03 |
| [axonogenesis](http://www.ebi.ac.uk/QuickGO/GTerm?id=GO:0007409" \o "http://www.ebi.ac.uk/QuickGO/GTerm?id=GO:0007409) | 1.00E-02 |
| [regulation of synaptic vesicle endocytosis](http://www.ebi.ac.uk/QuickGO/GTerm?id=GO:1900242" \o "http://www.ebi.ac.uk/QuickGO/GTerm?id=GO:1900242) | 1.10E-02 |
| [cellular response to interferon-gamma](http://www.ebi.ac.uk/QuickGO/GTerm?id=GO:0071346" \o "http://www.ebi.ac.uk/QuickGO/GTerm?id=GO:0071346) | 1.20E-02 |
| [receptor-mediated endocytosis](http://www.ebi.ac.uk/QuickGO/GTerm?id=GO:0006898" \o "http://www.ebi.ac.uk/QuickGO/GTerm?id=GO:0006898) | 1.20E-02 |
| [response to growth hormone](http://www.ebi.ac.uk/QuickGO/GTerm?id=GO:0060416" \o "http://www.ebi.ac.uk/QuickGO/GTerm?id=GO:0060416) | 1.20E-02 |
| [non-canonical Wnt signaling pathway](http://www.ebi.ac.uk/QuickGO/GTerm?id=GO:0035567" \o "http://www.ebi.ac.uk/QuickGO/GTerm?id=GO:0035567) | 1.20E-02 |
| [positive regulation of chemotaxis](http://www.ebi.ac.uk/QuickGO/GTerm?id=GO:0050921" \o "http://www.ebi.ac.uk/QuickGO/GTerm?id=GO:0050921) | 1.20E-02 |
| [response to peptide hormone](http://www.ebi.ac.uk/QuickGO/GTerm?id=GO:0043434" \o "http://www.ebi.ac.uk/QuickGO/GTerm?id=GO:0043434) | 1.30E-02 |
| [negative regulation of collateral sprouting of intact axon in response to injury](http://www.ebi.ac.uk/QuickGO/GTerm?id=GO:0048685" \o "http://www.ebi.ac.uk/QuickGO/GTerm?id=GO:0048685) | 1.40E-02 |
| [response to glucocorticoid](http://www.ebi.ac.uk/QuickGO/GTerm?id=GO:0051384" \o "http://www.ebi.ac.uk/QuickGO/GTerm?id=GO:0051384) | 1.40E-02 |
| [negative regulation of canonical Wnt signaling pathway](http://www.ebi.ac.uk/QuickGO/GTerm?id=GO:0090090" \o "http://www.ebi.ac.uk/QuickGO/GTerm?id=GO:0090090) | 1.50E-02 |
| [blood coagulation](http://www.ebi.ac.uk/QuickGO/GTerm?id=GO:0007596" \o "http://www.ebi.ac.uk/QuickGO/GTerm?id=GO:0007596) | 1.50E-02 |
| [circadian rhythm](http://www.ebi.ac.uk/QuickGO/GTerm?id=GO:0007623" \o "http://www.ebi.ac.uk/QuickGO/GTerm?id=GO:0007623) | 1.60E-02 |
| [cellular sodium ion homeostasis](http://www.ebi.ac.uk/QuickGO/GTerm?id=GO:0006883" \o "http://www.ebi.ac.uk/QuickGO/GTerm?id=GO:0006883) | 1.70E-02 |
| [response to estradiol](http://www.ebi.ac.uk/QuickGO/GTerm?id=GO:0032355" \o "http://www.ebi.ac.uk/QuickGO/GTerm?id=GO:0032355) | 1.80E-02 |
| [cellular response to cAMP](http://www.ebi.ac.uk/QuickGO/GTerm?id=GO:0071320" \o "http://www.ebi.ac.uk/QuickGO/GTerm?id=GO:0071320) | 1.80E-02 |
| [immunoglobulin mediated immune response](http://www.ebi.ac.uk/QuickGO/GTerm?id=GO:0016064" \o "http://www.ebi.ac.uk/QuickGO/GTerm?id=GO:0016064) | 2.00E-02 |
| [gene expression](http://www.ebi.ac.uk/QuickGO/GTerm?id=GO:0010467" \o "http://www.ebi.ac.uk/QuickGO/GTerm?id=GO:0010467) | 2.20E-02 |
| [response to oxidative stress](http://www.ebi.ac.uk/QuickGO/GTerm?id=GO:0006979" \o "http://www.ebi.ac.uk/QuickGO/GTerm?id=GO:0006979) | 2.30E-02 |
| [Wnt signaling pathway](http://www.ebi.ac.uk/QuickGO/GTerm?id=GO:0016055" \o "http://www.ebi.ac.uk/QuickGO/GTerm?id=GO:0016055) | 2.30E-02 |
| [lipid metabolic process](http://www.ebi.ac.uk/QuickGO/GTerm?id=GO:0006629" \o "http://www.ebi.ac.uk/QuickGO/GTerm?id=GO:0006629) | 2.30E-02 |
| [keratinocyte proliferation](http://www.ebi.ac.uk/QuickGO/GTerm?id=GO:0043616" \o "http://www.ebi.ac.uk/QuickGO/GTerm?id=GO:0043616) | 2.40E-02 |
| [negative regulation of proteolysis](http://www.ebi.ac.uk/QuickGO/GTerm?id=GO:0045861" \o "http://www.ebi.ac.uk/QuickGO/GTerm?id=GO:0045861) | 2.40E-02 |
| [cell-matrix adhesion](http://www.ebi.ac.uk/QuickGO/GTerm?id=GO:0007160" \o "http://www.ebi.ac.uk/QuickGO/GTerm?id=GO:0007160) | 2.60E-02 |
| [metanephric distal convoluted tubule development](http://www.ebi.ac.uk/QuickGO/GTerm?id=GO:0072221" \o "http://www.ebi.ac.uk/QuickGO/GTerm?id=GO:0072221) | 2.70E-02 |
| [Factor XII activation](http://www.ebi.ac.uk/QuickGO/GTerm?id=GO:0002542" \o "http://www.ebi.ac.uk/QuickGO/GTerm?id=GO:0002542) | 2.70E-02 |
| [negative regulation of extracellular matrix disassembly](http://www.ebi.ac.uk/QuickGO/GTerm?id=GO:0010716" \o "http://www.ebi.ac.uk/QuickGO/GTerm?id=GO:0010716) | 2.70E-02 |
| [organ or tissue specific immune response](http://www.ebi.ac.uk/QuickGO/GTerm?id=GO:0002251" \o "http://www.ebi.ac.uk/QuickGO/GTerm?id=GO:0002251) | 2.70E-02 |
| [creatinine homeostasis](http://www.ebi.ac.uk/QuickGO/GTerm?id=GO:0097273" \o "http://www.ebi.ac.uk/QuickGO/GTerm?id=GO:0097273) | 2.70E-02 |
| [cellular response to interleukin-6](http://www.ebi.ac.uk/QuickGO/GTerm?id=GO:0071354" \o "http://www.ebi.ac.uk/QuickGO/GTerm?id=GO:0071354) | 2.80E-02 |
| [positive regulation of gene expression](http://www.ebi.ac.uk/QuickGO/GTerm?id=GO:0010628" \o "http://www.ebi.ac.uk/QuickGO/GTerm?id=GO:0010628) | 2.80E-02 |
| [positive regulation of ERK1 and ERK2 cascade](http://www.ebi.ac.uk/QuickGO/GTerm?id=GO:0070374" \o "http://www.ebi.ac.uk/QuickGO/GTerm?id=GO:0070374) | 2.80E-02 |
| [decidualization](http://www.ebi.ac.uk/QuickGO/GTerm?id=GO:0046697" \o "http://www.ebi.ac.uk/QuickGO/GTerm?id=GO:0046697) | 2.90E-02 |
| [immune system process](http://www.ebi.ac.uk/QuickGO/GTerm?id=GO:0002376" \o "http://www.ebi.ac.uk/QuickGO/GTerm?id=GO:0002376) | 2.90E-02 |
| [male gonad development](http://www.ebi.ac.uk/QuickGO/GTerm?id=GO:0008584" \o "http://www.ebi.ac.uk/QuickGO/GTerm?id=GO:0008584) | 2.90E-02 |
| [androgen catabolic process](http://www.ebi.ac.uk/QuickGO/GTerm?id=GO:0006710" \o "http://www.ebi.ac.uk/QuickGO/GTerm?id=GO:0006710) | 3.40E-02 |
| [folic acid import into cell](http://www.ebi.ac.uk/QuickGO/GTerm?id=GO:1904447" \o "http://www.ebi.ac.uk/QuickGO/GTerm?id=GO:1904447) | 3.40E-02 |
| [negative regulation of mature B cell apoptotic process](http://www.ebi.ac.uk/QuickGO/GTerm?id=GO:0002906" \o "http://www.ebi.ac.uk/QuickGO/GTerm?id=GO:0002906) | 3.40E-02 |
| [extracellular matrix organization](http://www.ebi.ac.uk/QuickGO/GTerm?id=GO:0030198" \o "http://www.ebi.ac.uk/QuickGO/GTerm?id=GO:0030198) | 3.50E-02 |
| [substantia nigra development](http://www.ebi.ac.uk/QuickGO/GTerm?id=GO:0021762" \o "http://www.ebi.ac.uk/QuickGO/GTerm?id=GO:0021762) | 3.80E-02 |
| [cellular response to glucocorticoid stimulus](http://www.ebi.ac.uk/QuickGO/GTerm?id=GO:0071385" \o "http://www.ebi.ac.uk/QuickGO/GTerm?id=GO:0071385) | 3.80E-02 |
| [regulation of complement-dependent cytotoxicity](http://www.ebi.ac.uk/QuickGO/GTerm?id=GO:1903659" \o "http://www.ebi.ac.uk/QuickGO/GTerm?id=GO:1903659) | 4.00E-02 |
| [lipoprotein catabolic process](http://www.ebi.ac.uk/QuickGO/GTerm?id=GO:0042159" \o "http://www.ebi.ac.uk/QuickGO/GTerm?id=GO:0042159) | 4.00E-02 |
| [aflatoxin catabolic process](http://www.ebi.ac.uk/QuickGO/GTerm?id=GO:0046223" \o "http://www.ebi.ac.uk/QuickGO/GTerm?id=GO:0046223) | 4.00E-02 |
| [glutathione derivative biosynthetic process](http://www.ebi.ac.uk/QuickGO/GTerm?id=GO:1901687" \o "http://www.ebi.ac.uk/QuickGO/GTerm?id=GO:1901687) | 4.00E-02 |
| [regulation of microvillus length](http://www.ebi.ac.uk/QuickGO/GTerm?id=GO:0032532" \o "http://www.ebi.ac.uk/QuickGO/GTerm?id=GO:0032532) | 4.00E-02 |
| [regulation of neurotransmitter receptor activity](http://www.ebi.ac.uk/QuickGO/GTerm?id=GO:0099601" \o "http://www.ebi.ac.uk/QuickGO/GTerm?id=GO:0099601) | 4.00E-02 |
| [neuron projection development](http://www.ebi.ac.uk/QuickGO/GTerm?id=GO:0031175" \o "http://www.ebi.ac.uk/QuickGO/GTerm?id=GO:0031175) | 4.20E-02 |
| [positive regulation of axon extension](http://www.ebi.ac.uk/QuickGO/GTerm?id=GO:0045773" \o "http://www.ebi.ac.uk/QuickGO/GTerm?id=GO:0045773) | 4.20E-02 |
| [response to inorganic substance](http://www.ebi.ac.uk/QuickGO/GTerm?id=GO:0010035" \o "http://www.ebi.ac.uk/QuickGO/GTerm?id=GO:0010035) | 4.40E-02 |
| [defense response](http://www.ebi.ac.uk/QuickGO/GTerm?id=GO:0006952" \o "http://www.ebi.ac.uk/QuickGO/GTerm?id=GO:0006952) | 4.40E-02 |
| [response to lead ion](http://www.ebi.ac.uk/QuickGO/GTerm?id=GO:0010288" \o "http://www.ebi.ac.uk/QuickGO/GTerm?id=GO:0010288) | 4.50E-02 |
| [positive regulation of protein kinase B signaling](http://www.ebi.ac.uk/QuickGO/GTerm?id=GO:0051897" \o "http://www.ebi.ac.uk/QuickGO/GTerm?id=GO:0051897) | 4.60E-02 |
| [negative regulation of platelet activation](http://www.ebi.ac.uk/QuickGO/GTerm?id=GO:0010544" \o "http://www.ebi.ac.uk/QuickGO/GTerm?id=GO:0010544) | 4.70E-02 |
| [positive regulation of apoptotic cell clearance](http://www.ebi.ac.uk/QuickGO/GTerm?id=GO:2000427" \o "http://www.ebi.ac.uk/QuickGO/GTerm?id=GO:2000427) | 4.70E-02 |
| [vitamin D metabolic process](http://www.ebi.ac.uk/QuickGO/GTerm?id=GO:0042359" \o "http://www.ebi.ac.uk/QuickGO/GTerm?id=GO:0042359) | 4.70E-02 |
| [positive regulation of estradiol secretion](http://www.ebi.ac.uk/QuickGO/GTerm?id=GO:2000866" \o "http://www.ebi.ac.uk/QuickGO/GTerm?id=GO:2000866) | 4.70E-02 |
| [urate biosynthetic process](http://www.ebi.ac.uk/QuickGO/GTerm?id=GO:0034418" \o "http://www.ebi.ac.uk/QuickGO/GTerm?id=GO:0034418) | 4.70E-02 |
| [monocyte aggregation](http://www.ebi.ac.uk/QuickGO/GTerm?id=GO:0070487" \o "http://www.ebi.ac.uk/QuickGO/GTerm?id=GO:0070487) | 4.70E-02 |
| [response to activity](http://www.ebi.ac.uk/QuickGO/GTerm?id=GO:0014823" \o "http://www.ebi.ac.uk/QuickGO/GTerm?id=GO:0014823) | 4.70E-02 |
| [response to radiation](http://www.ebi.ac.uk/QuickGO/GTerm?id=GO:0009314" \o "http://www.ebi.ac.uk/QuickGO/GTerm?id=GO:0009314) | 4.90E-02 |
| [kidney development](http://www.ebi.ac.uk/QuickGO/GTerm?id=GO:0001822" \o "http://www.ebi.ac.uk/QuickGO/GTerm?id=GO:0001822) | 4.90E-02 |

Table S7 Signaling Pathways Enriched for Differential Proteins Produced under Relaxed Conditions Compared to Day0 vs. Day4 (P<0.05)

| [pathway](https://david.ncifcrf.gov/chartReport.jsp?d-16544-p=1&d-16544-o=2&annot=52&d-16544-s=2" \o "https://david.ncifcrf.gov/chartReport.jsp?d-16544-p=1&d-16544-o=2&annot=52&d-16544-s=2) | [P-Value](https://david.ncifcrf.gov/chartReport.jsp?d-16544-p=1&d-16544-o=1&annot=52&d-16544-s=7" \o "https://david.ncifcrf.gov/chartReport.jsp?d-16544-p=1&d-16544-o=1&annot=52&d-16544-s=7) |
| --- | --- |
| [Proteoglycans in cancer](https://david.ncifcrf.gov/kegg.jsp?path=rno05205$Proteoglycans in cancer&termId=520135532&source=kegg" \o "https://david.ncifcrf.gov/kegg.jsp?path=rno05205$Proteoglycans in cancer&termId=520135532&source=kegg) | 7.20E-07 |
| [Complement and coagulation cascades](https://david.ncifcrf.gov/kegg.jsp?path=rno04610$Complement and coagulation cascades&termId=520135405&source=kegg" \o "https://david.ncifcrf.gov/kegg.jsp?path=rno04610$Complement and coagulation cascades&termId=520135405&source=kegg) | 5.90E-06 |
| [Leukocyte transendothelial migration](https://david.ncifcrf.gov/kegg.jsp?path=rno04670$Leukocyte transendothelial migration&termId=520135426&source=kegg" \o "https://david.ncifcrf.gov/kegg.jsp?path=rno04670$Leukocyte transendothelial migration&termId=520135426&source=kegg) | 4.60E-05 |
| [Regulation of actin cytoskeleton](https://david.ncifcrf.gov/kegg.jsp?path=rno04810$Regulation of actin cytoskeleton&termId=520135445&source=kegg" \o "https://david.ncifcrf.gov/kegg.jsp?path=rno04810$Regulation of actin cytoskeleton&termId=520135445&source=kegg) | 1.00E-04 |
| [Protein digestion and absorption](https://david.ncifcrf.gov/kegg.jsp?path=rno04974$Protein digestion and absorption&termId=520135485&source=kegg" \o "https://david.ncifcrf.gov/kegg.jsp?path=rno04974$Protein digestion and absorption&termId=520135485&source=kegg) | 1.70E-04 |
| [Focal adhesion](https://david.ncifcrf.gov/kegg.jsp?path=rno04510$Focal adhesion&termId=520135398&source=kegg" \o "https://david.ncifcrf.gov/kegg.jsp?path=rno04510$Focal adhesion&termId=520135398&source=kegg) | 2.10E-04 |
| [Fluid shear stress and atherosclerosis](https://david.ncifcrf.gov/kegg.jsp?path=rno05418$Fluid shear stress and atherosclerosis&termId=520135570&source=kegg" \o "https://david.ncifcrf.gov/kegg.jsp?path=rno05418$Fluid shear stress and atherosclerosis&termId=520135570&source=kegg) | 2.10E-04 |
| [Gastric acid secretion](https://david.ncifcrf.gov/kegg.jsp?path=rno04971$Gastric acid secretion&termId=520135482&source=kegg" \o "https://david.ncifcrf.gov/kegg.jsp?path=rno04971$Gastric acid secretion&termId=520135482&source=kegg) | 3.50E-04 |
| [Hepatocellular carcinoma](https://david.ncifcrf.gov/kegg.jsp?path=rno05225$Hepatocellular carcinoma&termId=520135551&source=kegg" \o "https://david.ncifcrf.gov/kegg.jsp?path=rno05225$Hepatocellular carcinoma&termId=520135551&source=kegg) | 6.00E-04 |
| [ECM-receptor interaction](https://david.ncifcrf.gov/kegg.jsp?path=rno04512$ECM-receptor interaction&termId=520135399&source=kegg" \o "https://david.ncifcrf.gov/kegg.jsp?path=rno04512$ECM-receptor interaction&termId=520135399&source=kegg) | 7.00E-04 |
| [Hippo signaling pathway](https://david.ncifcrf.gov/kegg.jsp?path=rno04390$Hippo signaling pathway&termId=520135396&source=kegg" \o "https://david.ncifcrf.gov/kegg.jsp?path=rno04390$Hippo signaling pathway&termId=520135396&source=kegg) | 1.70E-03 |
| [Tight junction](https://david.ncifcrf.gov/kegg.jsp?path=rno04530$Tight junction&termId=520135402&source=kegg" \o "https://david.ncifcrf.gov/kegg.jsp?path=rno04530$Tight junction&termId=520135402&source=kegg) | 2.40E-03 |
| [Renin-angiotensin system](https://david.ncifcrf.gov/kegg.jsp?path=rno04614$Renin-angiotensin system&termId=520135409&source=kegg" \o "https://david.ncifcrf.gov/kegg.jsp?path=rno04614$Renin-angiotensin system&termId=520135409&source=kegg) | 2.50E-03 |
| [Glutathione metabolism](https://david.ncifcrf.gov/kegg.jsp?path=rno00480$Glutathione metabolism&termId=520135256&source=kegg" \o "https://david.ncifcrf.gov/kegg.jsp?path=rno00480$Glutathione metabolism&termId=520135256&source=kegg) | 2.70E-03 |
| [Bacterial invasion of epithelial cells](https://david.ncifcrf.gov/kegg.jsp?path=rno05100$Bacterial invasion of epithelial cells&termId=520135503&source=kegg" \o "https://david.ncifcrf.gov/kegg.jsp?path=rno05100$Bacterial invasion of epithelial cells&termId=520135503&source=kegg) | 3.60E-03 |
| [Pathways in cancer](https://david.ncifcrf.gov/kegg.jsp?path=rno05200$Pathways in cancer&termId=520135528&source=kegg" \o "https://david.ncifcrf.gov/kegg.jsp?path=rno05200$Pathways in cancer&termId=520135528&source=kegg) | 3.70E-03 |
| [Apoptosis](https://david.ncifcrf.gov/kegg.jsp?path=rno04210$Apoptosis&termId=520135378&source=kegg" \o "https://david.ncifcrf.gov/kegg.jsp?path=rno04210$Apoptosis&termId=520135378&source=kegg) | 4.90E-03 |
| [Hematopoietic cell lineage](https://david.ncifcrf.gov/kegg.jsp?path=rno04640$Hematopoietic cell lineage&termId=520135416&source=kegg" \o "https://david.ncifcrf.gov/kegg.jsp?path=rno04640$Hematopoietic cell lineage&termId=520135416&source=kegg) | 5.90E-03 |
| [Influenza A](https://david.ncifcrf.gov/kegg.jsp?path=rno05164$Influenza A&termId=520135520&source=kegg" \o "https://david.ncifcrf.gov/kegg.jsp?path=rno05164$Influenza A&termId=520135520&source=kegg) | 1.20E-02 |
| [Cell adhesion molecules](https://david.ncifcrf.gov/kegg.jsp?path=rno04514$Cell adhesion molecules&termId=520135400&source=kegg" \o "https://david.ncifcrf.gov/kegg.jsp?path=rno04514$Cell adhesion molecules&termId=520135400&source=kegg) | 1.50E-02 |
| [Phagosome](https://david.ncifcrf.gov/kegg.jsp?path=rno04145$Phagosome&termId=520135373&source=kegg" \o "https://david.ncifcrf.gov/kegg.jsp?path=rno04145$Phagosome&termId=520135373&source=kegg) | 1.70E-02 |
| [Platelet activation](https://david.ncifcrf.gov/kegg.jsp?path=rno04611$Platelet activation&termId=520135406&source=kegg" \o "https://david.ncifcrf.gov/kegg.jsp?path=rno04611$Platelet activation&termId=520135406&source=kegg) | 1.90E-02 |
| [Cytokine-cytokine receptor interaction](https://david.ncifcrf.gov/kegg.jsp?path=rno04060$Cytokine-cytokine receptor interaction&termId=520135351&source=kegg" \o "https://david.ncifcrf.gov/kegg.jsp?path=rno04060$Cytokine-cytokine receptor interaction&termId=520135351&source=kegg) | 2.20E-02 |
| [PI3K-Akt signaling pathway](https://david.ncifcrf.gov/kegg.jsp?path=rno04151$PI3K-Akt signaling pathway&termId=520135376&source=kegg" \o "https://david.ncifcrf.gov/kegg.jsp?path=rno04151$PI3K-Akt signaling pathway&termId=520135376&source=kegg) | 2.20E-02 |
| [Human papillomavirus infection](https://david.ncifcrf.gov/kegg.jsp?path=rno05165$Human papillomavirus infection&termId=520135521&source=kegg" \o "https://david.ncifcrf.gov/kegg.jsp?path=rno05165$Human papillomavirus infection&termId=520135521&source=kegg) | 2.30E-02 |
| [Yersinia infection](https://david.ncifcrf.gov/kegg.jsp?path=rno05135$Yersinia infection&termId=520135507&source=kegg" \o "https://david.ncifcrf.gov/kegg.jsp?path=rno05135$Yersinia infection&termId=520135507&source=kegg) | 2.40E-02 |
| [Arrhythmogenic right ventricular cardiomyopathy](https://david.ncifcrf.gov/kegg.jsp?path=rno05412$Arrhythmogenic right ventricular cardiomyopathy&termId=520135565&source=kegg" \o "https://david.ncifcrf.gov/kegg.jsp?path=rno05412$Arrhythmogenic right ventricular cardiomyopathy&termId=520135565&source=kegg) | 2.50E-02 |
| [Rap1 signaling pathway](https://david.ncifcrf.gov/kegg.jsp?path=rno04015$Rap1 signaling pathway&termId=520135347&source=kegg" \o "https://david.ncifcrf.gov/kegg.jsp?path=rno04015$Rap1 signaling pathway&termId=520135347&source=kegg) | 2.90E-02 |
| [Viral myocarditis](https://david.ncifcrf.gov/kegg.jsp?path=rno05416$Viral myocarditis&termId=520135568&source=kegg" \o "https://david.ncifcrf.gov/kegg.jsp?path=rno05416$Viral myocarditis&termId=520135568&source=kegg) | 3.20E-02 |
| [Oxytocin signaling pathway](https://david.ncifcrf.gov/kegg.jsp?path=rno04921$Oxytocin signaling pathway&termId=520135458&source=kegg" \o "https://david.ncifcrf.gov/kegg.jsp?path=rno04921$Oxytocin signaling pathway&termId=520135458&source=kegg) | 3.60E-02 |
| [Hypertrophic cardiomyopathy](https://david.ncifcrf.gov/kegg.jsp?path=rno05410$Hypertrophic cardiomyopathy&termId=520135564&source=kegg" \o "https://david.ncifcrf.gov/kegg.jsp?path=rno05410$Hypertrophic cardiomyopathy&termId=520135564&source=kegg) | 3.80E-02 |
| [Adherens junction](https://david.ncifcrf.gov/kegg.jsp?path=rno04520$Adherens junction&termId=520135401&source=kegg" \o "https://david.ncifcrf.gov/kegg.jsp?path=rno04520$Adherens junction&termId=520135401&source=kegg) | 4.00E-02 |
| [Dilated cardiomyopathy](https://david.ncifcrf.gov/kegg.jsp?path=rno05414$Dilated cardiomyopathy&termId=520135566&source=kegg" \o "https://david.ncifcrf.gov/kegg.jsp?path=rno05414$Dilated cardiomyopathy&termId=520135566&source=kegg) | 4.10E-02 |
| [Staphylococcus aureus infection](https://david.ncifcrf.gov/kegg.jsp?path=rno05150$Staphylococcus aureus infection&termId=520135514&source=kegg" \o "https://david.ncifcrf.gov/kegg.jsp?path=rno05150$Staphylococcus aureus infection&termId=520135514&source=kegg) | 4.70E-02 |

Table S8 Differential protein production by rats sniffing anemones (FC≥1.5 or ≤0.67, P<0.05)

| Accession | Protein names | Day10 vs Day13 | | | | Day10 vs Day14 | | |
| --- | --- | --- | --- | --- | --- | --- | --- | --- |
|  |  | Trend | Fold change | | P value | Trend | Fold change | P value |
| A0A0G2JWX4 | Keratin 2 | ↓ | 0.24 | 2.84E-02 | | ↓ | 0.25 | 2.91E-02 |
| A0A0G2K2V6 | Keratin 10 | ↓ | 0.33 | 3.48E-02 | | ↓ | 0.31 | 3.04E-02 |
| Q4FZU2 | Keratin, type II cytoskeletal 6A | ↓ | 0.34 | 2.97E-02 | | ↓ | 0.32 | 2.69E-02 |
| P29975 | Aquaporin-1 | ↓ | 0.37 | 1.24E-04 | | ↓ | 0.56 | 6.51E-03 |
| P17559 | Uteroglobin | ↓ | 0.38 | 1.60E-03 | | - | - | - |
| M0R9A3 | PBPC1BS-like | ↓ | 0.41 | 2.01E-02 | | - | - | - |
| B2GV31 | Muc1 protein | ↓ | 0.41 | 1.37E-04 | | ↓ | 0.47 | 3.14E-04 |
| Q6P6Q2 | Keratin, type II cytoskeletal 5 | ↓ | 0.42 | 3.45E-02 | | - | - | - |
| Q6IFV1 | Keratin, type I cytoskeletal 14 | ↓ | 0.43 | 4.58E-02 | | ↓ | 0.40 | 3.01E-02 |
| A0A0G2JST3 | Keratin, type II cytoskeletal 1 | ↓ | 0.45 | 4.30E-02 | | ↓ | 0.39 | 2.59E-02 |
| B1WC34 | Protein kinase C substrate 80K-H | ↓ | 0.46 | 9.08E-05 | | ↓ | 0.39 | 2.16E-05 |
| M0RCH6 | Charged multivesicular body protein 4B | ↓ | 0.47 | 2.02E-03 | | ↓ | 0.43 | 9.94E-04 |
| Q9QZH0 | Frizzled-4 | ↓ | 0.48 | 1.43E-03 | | ↓ | 0.35 | 3.16E-04 |
| A0A0G2JXL5 | Mucin 19, oligomeric | ↓ | 0.50 | 4.73E-03 | | ↓ | 0.45 | 2.48E-03 |
| G3V7Y3 | ATP synthase F1 subunit delta | ↓ | 0.50 | 9.23E-05 | | - | - | - |
| F1LRT1 | fructose-bisphosphatase | ↓ | 0.55 | 5.22E-04 | | - | - | - |
| A0A0G2K1F0 | Ig-like domain-containing protein | ↓ | 0.56 | 7.81E-03 | | ↓ | 0.54 | 2.80E-03 |
| P81827 | Urinary protein 1 | ↓ | 0.57 | 4.53E-03 | | ↓ | 0.50 | 3.31E-03 |
| D3ZHD1 | Annexin | ↓ | 0.58 | 2.23E-02 | | - | - | - |
| P51146 | Ras-related protein Rab-4B | ↓ | 0.58 | 2.71E-03 | | ↓ | 0.60 | 8.72E-03 |
| Q80W57 | Broad substrate specificity ATP-binding cassette transporter ABCG2 | ↓ | 0.59 | 3.88E-02 | | - | - | - |
| Q8CG08 | Collagen triple helix repeat-containing protein 1 | ↓ | 0.59 | 1.10E-02 | | ↓ | 0.47 | 1.42E-03 |
| Q6TMA8 | Angiopoietin-related protein 4 | ↓ | 0.59 | 2.53E-02 | | ↓ | 0.46 | 1.26E-03 |
| P02454 | Collagen alpha-1 | ↓ | 0.59 | 2.93E-02 | | ↓ | 0.62 | 4.66E-02 |
| B2RZB5 | Charged multivesicular body protein 2A | ↓ | 0.60 | 8.57E-04 | | ↓ | 0.51 | 3.43E-04 |
| P55159 | Serum paraoxonase/arylesterase 1 | ↓ | 0.60 | 1.01E-02 | | - | - | - |
| G3V8X4 | Gliomedin | ↓ | 0.60 | 3.52E-02 | | ↓ | 0.49 | 6.44E-03 |
| Q5BJY9 | Keratin, type I cytoskeletal 18 | ↓ | 0.60 | 4.94E-02 | | - | - | - |
| D3ZN06 | CD248 molecule | ↓ | 0.60 | 3.11E-03 | | - | - | - |
| P07335 | Creatine kinase B-type | ↓ | 0.61 | 4.45E-02 | | - | - | - |
| A0A0G2QC46 | Roundabout guidance receptor 4 | ↓ | 0.61 | 1.57E-02 | | ↓ | 0.58 | 1.76E-02 |
| D3ZLA3 | Copine 3 | ↓ | 0.62 | 1.28E-03 | | - | - | - |
| A0A0G2K4K4 | Solute carrier family 12 member 1 | ↓ | 0.63 | 2.10E-02 | | - | - | - |
| Q568Z6 | IST1 homolog | ↓ | 0.63 | 2.32E-03 | | ↓ | 0.65 | 8.23E-03 |
| D3ZUK3 | Protein delta homolog 2 | ↓ | 0.65 | 3.23E-02 | | ↓ | 0.58 | 7.46E-03 |
| A0A0G2JTH4 | Leukocyte surface antigen CD47 | ↓ | 0.65 | 8.43E-03 | | - | - | - |
| A4KWA5 | C-type lectin domain family 2 member D2 | ↓ | 0.65 | 8.42E-03 | | - | - | - |
| P19939 | Apolipoprotein C-I | ↓ | 0.66 | 2.29E-02 | | ↓ | 0.57 | 8.94E-03 |
| Q4V8K5 | BRO1 domain-containing protein BROX | ↓ | 0.66 | 3.75E-03 | | - | - | - |
| B0BND0 | Glycerophosphocholine cholinephosphodiesterase ENPP6 | ↓ | 0.66 | 8.49E-04 | | ↓ | 0.67 | 2.14E-03 |
| A0A0G2K2P7 | Serine/threonine-protein kinase receptor | ↓ | 0.66 | 7.97E-03 | | ↓ | 0.52 | 2.29E-03 |
| G3V7X5 | SPARC-like protein 1 | ↓ | 0.66 | 4.71E-02 | | - | - | - |
| A0A0H2UHE4 | Regenerating family member 3 beta | ↓ | 0.66 | 9.08E-03 | | - | - | - |
| D3ZUU6 | C-type lectin domain family 3, member B | ↓ | 0.67 | 2.31E-02 | | - | - | - |
| F1LRE2 | Insulin-like growth factor binding protein,IGFBP2 | ↑ | 1.54 | 3.07E-02 | | ↑ | 2.08 | 1.31E-03 |
| A0A0G2JSR9 | 45 kDa calcium-binding protein | ↑ | 1.54 | 4.92E-02 | | - | - | - |
| Q9R1T1 | Barrier-to-autointegration factor | ↑ | 1.54 | 5.40E-03 | | - | - | - |
| F1LQ83 | Achaete-scute family bHLH transcription factor 3 | ↑ | 1.55 | 1.66E-02 | | - | - | - |
| D4A281 | TNF receptor superfamily member 13C | ↑ | 1.57 | 1.12E-03 | | ↑ | 1.72 | 2.09E-05 |
| D3ZK14 | Tenascin N | ↑ | 1.57 | 4.64E-02 | | ↑ | 1.77 | 4.59E-03 |
| A0A0G2JVN2 | Secreted frizzled-related protein 4 | ↑ | 1.60 | 6.84E-03 | | - | - | - |
| Q5U300 | Ubiquitin-like modifier-activating enzyme 1 | ↑ | 1.77 | 1.93E-02 | | - | - | - |
| P35213 | 14-3-3 protein beta/alpha | ↑ | 1.78 | 6.21E-03 | | - | - | - |
| A0A0G2JSZ3 | Neuroblastoma suppressor of tumorigenicity 1 | ↑ | 1.86 | 1.22E-03 | | ↑ | 2.28 | 9.38E-06 |
| P35053 | Glypican-1 | ↑ | 2.12 | 2.90E-02 | | ↑ | 2.29 | 3.17E-02 |
| A0A0G2JVP4 | Immunoglobulin heavy constant epsilon | ↑ | 2.27 | 6.83E-03 | | ↑ | 1.74 | 4.28E-02 |
| G3V9J1 | Murinoglobulin 2 | ↑ | 2.31 | 2.58E-04 | | ↑ | 2.11 | 3.41E-03 |
| P16636 | Protein-lysine 6-oxidase | ↑ | 2.33 | 8.54E-04 | | ↑ | 2.93 | 3.34E-03 |
| A0A0G2JSY6 | Trefoil factor 3 | ↑ | 2.65 | 7.19E-05 | | ↑ | 2.77 | 4.79E-05 |
| A0A096MJ68 | Secretory leukocyte peptidase inhibitor | ↑ | 2.82 | 2.25E-03 | | ↑ | 2.87 | 2.28E-02 |
| P15399 | Probasin | ↑ | 2.94 | 1.31E-02 | | - | - | - |
| Q99MH3 | Hepcidin | ↑ | 3.08 | 8.45E-03 | | ↑ | 4.29 | 3.75E-05 |
| Q30KJ2 | Beta-defensin 50 | ↑ | 3.21 | 1.64E-02 | | - | - | - |
| O89117 | Beta-defensin 1 | ↑ | 3.49 | 1.22E-04 | | ↑ | 3.24 | 5.96E-03 |
| P97580 | Beta-microseminoprotein | ↑ | 3.66 | 3.24E-02 | | - | - | - |
| Q642B0 | Glypican 4 | ↑ | 4.18 | 1.34E-02 | | ↑ | 3.94 | 1.90E-02 |
| P07522 | Pro-epidermal growth factor |  |  |  | | ↓ | 0.40 | 9.67E-04 |
| Q6RUV5 | Ras-related C3 botulinum toxin substrate 1 |  |  |  | | ↓ | 0.41 | 2.39E-02 |
| P07756 | Carbamoyl-phosphate synthase |  |  |  | | ↓ | 0.44 | 2.45E-02 |
| P61589 | Transforming protein RhoA |  |  |  | | ↓ | 0.48 | 2.20E-02 |
| P46844 | Biliverdin reductase A |  |  |  | | ↓ | 0.50 | 1.83E-04 |
| Q9JJ19 | Na(+)/H(+) exchange regulatory cofactor NHE-RF1 |  |  |  | | ↓ | 0.50 | 3.53E-03 |
| F1LR02 | Collagen type XVIII alpha 1 chain |  |  |  | | ↓ | 0.51 | 3.18E-02 |
| P28902 | Guanylin |  |  |  | | ↓ | 0.53 | 4.40E-04 |
| Q63772 | Growth arrest-specific protein 6 |  |  |  | | ↓ | 0.55 | 4.55E-02 |
| Q5M843 | 2-oxoglutarate and iron-dependent oxygenase domain-containing protein 3 |  |  |  | | ↓ | 0.56 | 5.62E-05 |
| A0A0G2JZ18 | Pentraxin 4 |  |  |  | | ↓ | 0.58 | 1.61E-02 |
| P18418 | Calreticulin |  |  |  | | ↓ | 0.59 | 2.36E-02 |
| A0A0G2JZR4 | RAB11B, member RAS oncogene family |  |  |  | | ↓ | 0.60 | 1.21E-02 |
| P48508 | Glutamate--cysteine ligase regulatory subunit |  |  |  | | ↓ | 0.61 | 1.12E-02 |
| A0A0G2JU25 | Polypeptide N-acetylgalactosaminyltransferase |  |  |  | | ↓ | 0.61 | 1.80E-02 |
| O55145 | Fractalkine |  |  |  | | ↓ | 0.63 | 1.35E-02 |
| G3V989 | Ephrin A5 |  |  |  | | ↓ | 0.65 | 2.31E-02 |
| P62260 | 14-3-3 protein epsilon |  |  |  | | ↓ | 0.66 | 8.26E-03 |
| P61107 | Ras-related protein Rab-14 |  |  |  | | ↓ | 0.66 | 4.15E-03 |
| D4A5C0 | Nectin cell adhesion molecule 3 |  |  |  | | ↓ | 0.66 | 2.56E-02 |
| Q5XIE8 | Integral membrane protein 2B |  |  |  | | ↓ | 0.67 | 2.15E-03 |
| Q9Z339 | Glutathione S-transferase omega-1 |  |  |  | | ↓ | 0.67 | 2.87E-03 |
| P07483 | Fatty acid-binding protein, heart |  |  |  | | ↑ | 1.52 | 3.35E-02 |
| Q8CHN3 | WAP four-disulfide core domain protein 2 |  |  |  | | ↑ | 1.53 | 5.32E-03 |
| A0A0G2JSS8 | Peroxiredoxin-5 |  |  |  | | ↑ | 1.53 | 2.49E-02 |
| P07151 | Beta-2-microglobulin |  |  |  | | ↑ | 1.53 | 6.84E-03 |
| Q9R066 | Coxsackievirus and adenovirus receptor homolog |  |  |  | | ↑ | 1.55 | 3.15E-02 |
| D3ZVB7 | Osteoglycin |  |  |  | | ↑ | 1.56 | 1.24E-02 |
| P47853 | Biglycan |  |  |  | | ↑ | 1.57 | 1.86E-02 |
| P14844 | C-C motif chemokine 2 |  |  |  | | ↑ | 1.57 | 2.15E-02 |
| P50137 | Transketolase |  |  |  | | ↑ | 1.57 | 4.23E-02 |
| P15473 | Insulin-like growth factor-binding protein 3 |  |  |  | | ↑ | 1.58 | 2.19E-03 |
| G3V8V1 | Granulin precursor |  |  |  | | ↑ | 1.59 | 2.84E-03 |
| P35859 | Insulin-like growth factor-binding protein complex acid labile subunit |  |  |  | | ↑ | 1.60 | 7.25E-03 |
| P08649 | Complement C4 |  |  |  | | ↑ | 1.63 | 8.99E-04 |
| A0A0G2K013 | Actinin alpha 4 |  |  |  | | ↑ | 1.64 | 2.09E-02 |
| P32755 | 4-hydroxyphenylpyruvate dioxygenase |  |  |  | | ↑ | 1.71 | 7.66E-03 |
| A0A0H2UHR7 | Filamin-C |  |  |  | | ↑ | 1.73 | 5.54E-03 |
| D3ZSC1 | Sushi domain containing 5 |  |  |  | | ↑ | 1.73 | 6.24E-03 |
| D3ZTN6 | SREBP regulating gene protein |  |  |  | | ↑ | 1.74 | 2.84E-02 |
| P51635 | Aldo-keto reductase family 1 member A1 |  |  |  | | ↑ | 1.74 | 8.75E-03 |
| P20761 | Ig gamma-2B chain C region |  |  |  | | ↑ | 1.75 | 1.50E-02 |
| B3EY86 | Lipocalin 11 |  |  |  | | ↑ | 1.77 | 3.10E-02 |
| P0DMW0 | Heat shock 70 kDa protein 1A |  |  |  | | ↑ | 1.85 | 3.18E-02 |
| D3ZA76 | Serine protease HTRA3 |  |  |  | | ↑ | 1.92 | 1.69E-03 |
| D3ZJV7 | Lipocalin 4 |  |  |  | | ↑ | 1.93 | 1.64E-02 |
| Q5M890 | Apolipoprotein N |  |  |  | | ↑ | 2.06 | 1.56E-02 |
| P02631 | Oncomodulin |  |  |  | | ↑ | 2.19 | 8.72E-03 |
| A0A0G2JV31 | X-prolyl aminopeptidase 1 |  |  |  | | ↑ | 2.22 | 2.23E-02 |
| P27590 | Uromodulin |  |  |  | | ↑ | 2.24 | 2.81E-02 |
| D4A5L9 | Similar to Cytochrome c, somatic |  |  |  | | ↑ | 2.24 | 4.69E-03 |
| M0R5T8 | Ferritin |  |  |  | | ↑ | 2.45 | 7.44E-03 |

Table S9 Biological Processes Produced by Differential Protein Enrichment to Day10 vs. Day13 Under Relaxed Conditions (P<0.05)

| Biological process | P-Value |
| --- | --- |
| intermediate filament organization | 1.50E-06 |
| wound healing | 1.00E-04 |
| response to nutrient levels | 3.30E-04 |
| keratinization | 1.40E-03 |
| cell migration | 2.60E-03 |
| peptide cross-linking | 3.00E-03 |
| response to peptide hormone | 8.40E-03 |
| negative regulation of intestinal absorption | 9.30E-03 |
| negative regulation of inflammatory response | 1.10E-02 |
| positive regulation of cell migration | 1.20E-02 |
| positive regulation of epidermis development | 1.20E-02 |
| mitotic nuclear envelope reassembly | 1.50E-02 |
| regulation of protein localization to membrane | 1.50E-02 |
| regulation of bone development | 1.50E-02 |
| negative regulation of lipoprotein lipase activity | 1.90E-02 |
| cellular response to tumor necrosis factor | 2.30E-02 |
| cellular response to dexamethasone stimulus | 2.40E-02 |
| blood vessel development | 2.60E-02 |
| defense response to bacterium | 2.60E-02 |
| cellular response to retinoic acid | 2.80E-02 |
| positive regulation of fibroblast proliferation | 3.00E-02 |
| response to fluoride | 3.10E-02 |
| endothelial cell apoptotic process | 3.40E-02 |
| defense response to Gram-negative bacterium | 3.90E-02 |
| cellular response to phorbol 13-acetate 12-myristate | 4.00E-02 |
| antimicrobial humoral immune response mediated by antimicrobial peptide | 4.50E-02 |
| keratinocyte development | 4.60E-02 |
| osteoblast differentiation | 4.60E-02 |
| protein heterotetramerization | 4.90E-02 |

Table S10 Signaling Pathways Enriched for Differential Proteins Generated by Day10 vs. Day13 under Relaxed Conditions (P<0.05)

| pathway | P-Value |
| --- | --- |
| Staphylococcus aureus infection | 3.30E-03 |
| ECM-receptor interaction | 2.90E-02 |
| TGF-beta signaling pathway | 4.30E-02 |

Table S11 Biological Processes Produced by Differential Protein Enrichment to Day10 vs. Day14 Under Relaxed Conditions (P<0.05)

| Biological process | P-Value |
| --- | --- |
| positive regulation of cell migration | 4.40E-07 |
| response to xenobiotic stimulus | 7.70E-07 |
| wound healing | 4.60E-05 |
| response to hypoxia | 6.10E-05 |
| cortical cytoskeleton organization | 7.10E-05 |
| negative regulation of cell-substrate adhesion | 7.10E-05 |
| actin cytoskeleton organization | 7.10E-05 |
| substrate adhesion-dependent cell spreading | 1.30E-04 |
| intermediate filament organization | 1.60E-04 |
| negative regulation of intracellular steroid hormone receptor signaling pathway | 1.80E-04 |
| positive regulation of endothelial cell migration | 2.10E-04 |
| cell migration | 3.50E-04 |
| regulation of cell shape | 4.50E-04 |
| neuron projection morphogenesis | 5.40E-04 |
| neuron migration | 5.70E-04 |
| regulation of actin cytoskeleton organization | 9.20E-04 |
| hyperosmotic response | 9.80E-04 |
| cellular response to fibroblast growth factor stimulus | 1.20E-03 |
| negative regulation of fibroblast migration | 1.20E-03 |
| cell adhesion | 1.20E-03 |
| regulation of JAK-STAT cascade | 1.60E-03 |
| establishment or maintenance of cell polarity | 1.70E-03 |
| positive regulation of actin filament polymerization | 1.80E-03 |
| Wnt signaling pathway, planar cell polarity pathway | 2.10E-03 |
| response to mechanical stimulus | 2.20E-03 |
| regulation of cell migration | 2.20E-03 |
| positive regulation of stress fiber assembly | 2.30E-03 |
| cytoskeleton organization | 2.50E-03 |
| response to amino acid | 2.70E-03 |
| innate immune response | 2.70E-03 |
| cellular response to chemokine | 3.00E-03 |
| establishment of epithelial cell apical/basal polarity | 3.00E-03 |
| phagocytosis, engulfment | 3.30E-03 |
| positive regulation of cell adhesion | 3.30E-03 |
| negative regulation of inflammatory response | 3.80E-03 |
| aging | 3.90E-03 |
| positive regulation of NIK/NF-kappaB signaling | 4.70E-03 |
| non-canonical Wnt signaling pathway | 5.20E-03 |
| cellular response to retinoic acid | 5.50E-03 |
| peptide cross-linking | 5.60E-03 |
| positive regulation of fibroblast proliferation | 6.10E-03 |
| regulation of cell morphogenesis | 6.40E-03 |
| cochlea morphogenesis | 6.90E-03 |
| regulation of cell size | 6.90E-03 |
| negative regulation of neuron projection development | 7.10E-03 |
| cell-cell junction organization | 7.90E-03 |
| muscle cell development | 8.40E-03 |
| trabecula morphogenesis | 8.60E-03 |
| alpha-beta T cell lineage commitment | 8.60E-03 |
| anatomical structure arrangement | 8.60E-03 |
| skeletal muscle satellite cell migration | 8.60E-03 |
| cerebral cortex GABAergic interneuron development | 8.60E-03 |
| regulation of cell adhesion involved in heart morphogenesis | 8.60E-03 |
| mitotic cleavage furrow formation | 8.60E-03 |
| positive regulation of lamellipodium assembly | 8.90E-03 |
| small GTPase mediated signal transduction | 9.50E-03 |
| cellular response to tumor necrosis factor | 1.00E-02 |
| cellular response to lipopolysaccharide | 1.10E-02 |
| osteoblast differentiation | 1.20E-02 |
| defense response to bacterium | 1.20E-02 |
| regulation of microtubule cytoskeleton organization | 1.30E-02 |
| positive regulation of lipase activity | 1.30E-02 |
| embryonic olfactory bulb interneuron precursor migration | 1.30E-02 |
| endothelial tube lumen extension | 1.30E-02 |
| negative regulation of interleukin-23 production | 1.30E-02 |
| positive regulation of DNA replication | 1.30E-02 |
| positive regulation of dendritic spine development | 1.30E-02 |
| negative regulation of neuron apoptotic process | 1.40E-02 |
| positive regulation of neuron differentiation | 1.40E-02 |
| cellular response to interleukin-1 | 1.40E-02 |
| negative chemotaxis | 1.50E-02 |
| heterophilic cell-cell adhesion via plasma membrane cell adhesion molecules | 1.50E-02 |
| positive regulation of substrate adhesion-dependent cell spreading | 1.50E-02 |
| regulation of osteoblast proliferation | 1.70E-02 |
| positive regulation of epidermis development | 1.70E-02 |
| angiotensin-mediated vasoconstriction involved in regulation of systemic arterial blood pressure | 1.70E-02 |
| positive regulation of vascular smooth muscle contraction | 1.70E-02 |
| regulation of respiratory burst | 1.70E-02 |
| beta selection | 1.70E-02 |
| angiogenesis | 1.80E-02 |
| regulation of growth | 1.80E-02 |
| response to ethanol | 2.00E-02 |
| tumor necrosis factor-mediated signaling pathway | 2.00E-02 |
| response to glucocorticoid | 2.10E-02 |
| signal transduction | 2.10E-02 |
| regulation of protein localization to membrane | 2.10E-02 |
| localization within membrane | 2.10E-02 |
| establishment or maintenance of actin cytoskeleton polarity | 2.10E-02 |
| L-ascorbic acid biosynthetic process | 2.10E-02 |
| regulation of neuron maturation | 2.10E-02 |
| regulation of bone development | 2.10E-02 |
| cellular response to mechanical stimulus | 2.20E-02 |
| response to zinc ion | 2.30E-02 |
| protein localization to nucleus | 2.50E-02 |
| odontogenesis | 2.50E-02 |
| regulation of systemic arterial blood pressure by endothelin | 2.60E-02 |
| regulation of neutrophil migration | 2.60E-02 |
| mast cell chemotaxis | 2.60E-02 |
| positive regulation of ovarian follicle development | 2.60E-02 |
| negative regulation of lipoprotein lipase activity | 2.60E-02 |
| collagen fibril organization | 2.60E-02 |
| antibacterial humoral response | 2.90E-02 |
| positive regulation of apoptotic cell clearance | 3.00E-02 |
| auditory receptor cell morphogenesis | 3.00E-02 |
| apolipoprotein A-I-mediated signaling pathway | 3.00E-02 |
| regulation of modification of synaptic structure | 3.00E-02 |
| Wnt signaling pathway | 3.10E-02 |
| lipid metabolic process | 3.10E-02 |
| complement activation, classical pathway | 3.10E-02 |
| positive regulation of cysteine-type endopeptidase activity involved in apoptotic process | 3.10E-02 |
| response to steroid hormone | 3.10E-02 |
| negative regulation of cysteine-type endopeptidase activity involved in apoptotic process | 3.10E-02 |
| actin cytoskeleton reorganization | 3.30E-02 |
| Roundabout signaling pathway | 3.40E-02 |
| positive regulation of dendritic cell chemotaxis | 3.40E-02 |
| cleavage furrow formation | 3.40E-02 |
| negative regulation of microglial cell activation | 3.40E-02 |
| chemotaxis | 3.80E-02 |
| keratinization | 3.80E-02 |
| actin filament organization | 3.80E-02 |
| response to lipopolysaccharide | 3.80E-02 |
| apical junction assembly | 3.80E-02 |
| midbrain dopaminergic neuron differentiation | 3.80E-02 |
| regulation of actin polymerization or depolymerization | 3.80E-02 |
| water homeostasis | 3.80E-02 |
| negative regulation of oxidative phosphorylation | 3.80E-02 |
| negative regulation of cell migration | 4.00E-02 |
| protein localization to plasma membrane | 4.10E-02 |
| positive regulation of collateral sprouting | 4.20E-02 |
| forebrain radial glial cell differentiation | 4.20E-02 |
| regulation of protein localization to cell surface | 4.20E-02 |
| cell junction assembly | 4.20E-02 |
| cellular response to progesterone stimulus | 4.20E-02 |
| neutrophil chemotaxis | 4.30E-02 |
| cellular response to dexamethasone stimulus | 4.40E-02 |
| regulation of modification of postsynaptic actin cytoskeleton | 4.60E-02 |
| hepatocyte growth factor receptor signaling pathway | 4.60E-02 |
| regulation of lamellipodium assembly | 4.60E-02 |
| endothelial cell apoptotic process | 4.60E-02 |
| positive regulation of alpha-beta T cell differentiation | 4.60E-02 |
| blood vessel development | 4.80E-02 |
| negative regulation of neuron differentiation | 4.80E-02 |

Table S12 Signaling Pathways Enriched for Differential Proteins Produced by Relaxed Conditions Compared to Day10 vs. Day14 (P<0.05)

| pathway | P-Value |
| --- | --- |
| Focal adhesion | 7.00E-04 |
| Fluid shear stress and atherosclerosis | 1.30E-03 |
| Proteoglycans in cancer | 4.90E-03 |
| Human cytomegalovirus infection | 1.00E-02 |
| Adherens junction | 1.40E-02 |
| Staphylococcus aureus infection | 1.60E-02 |
| TGF-beta signaling pathway | 2.10E-02 |
| PI3K-Akt signaling pathway | 3.90E-02 |

Table S13 Differential proteins co-identified by 5 or more rats in the anemone group before and after the 6 rats themselves were compared (FC≥1.5 or ≤0.67, P<0.05)

| Accession | Protein name | Day0 vs.Day3 | Day0 vs.Day4 |
| --- | --- | --- | --- |
| Q99MH3 | Hepcidin | Rat1 Rat2 Rat3 Rat4 Rat5 Rat6 | Rat1 Rat3 Rat4 Rat5 Rat6 |
| F7EZQ4 | Fc fragment of IgG binding protein-like 1 | Rat1 Rat2 Rat3 Rat4 Rat5 Rat6 | Rat1 Rat2 Rat3 Rat4 Rat5 Rat6 |
| A0A0G2JSY6 | Trefoil factor 3 | Rat1 Rat2 Rat3 Rat4 Rat5 Rat6 | Rat1 Rat2 Rat3 Rat4 Rat5 Rat6 |
| Q5M890 | Apolipoprotein N | Rat1 Rat2 Rat3 Rat4 Rat5 Rat6 | Rat1 Rat2 Rat3 Rat4 Rat6 |
| Q8CG08 | Collagen triple helix repeat-containing protein 1 | Rat1 Rat2 Rat3 Rat4 Rat5 | Rat1 Rat2 Rat3 Rat4 Rat5 Rat6 |
| Q99041 | Protein-glutamine gamma-glutamyltransferase 4 | Rat1 Rat2 Rat3 Rat4 Rat5 | Rat1 Rat2 Rat3 Rat4 Rat5 Rat6 |
| B1WC34 | Protein kinase C substrate 80K-H | Rat1 Rat2 Rat3 Rat4 Rat6 | Rat1 Rat2 Rat3 Rat4 Rat6 |
| D3ZJV7 | Lipocalin 4 | Rat1 Rat2 Rat3 Rat4 Rat6 | Rat2 Rat3 Rat4 Rat5 Rat6 |
| Q9QZK9 | Deoxyribonuclease-2-beta | Rat1 Rat2 Rat3 Rat4 Rat6 | Rat1 Rat2 Rat3 Rat4 Rat6 |
| M0R6C7 | Mucin 2 | Rat1 Rat2 Rat3 Rat5 Rat6 | Rat1 Rat2 Rat3 Rat5 Rat6 |
| P07647 | Submandibular glandular kallikrein-9 | Rat1 Rat2 Rat3 Rat5 Rat6 | Rat1 Rat2 Rat3 Rat4 Rat6 |
| P97580 | Beta-microseminoprotein | Rat1 Rat2 Rat3 Rat5 Rat6 | Rat1 Rat2 Rat3 Rat4 Rat5 Rat6 |
| F1MAE5 | Proline rich protein 2-like 1 | Rat1 Rat2 Rat3 Rat5 Rat6 | Rat1 Rat2 Rat3 Rat4 Rat6 |
| Z4YNX7 | Cystatin-related protein 2 | Rat1 Rat2 Rat3 Rat5 Rat6 | Rat1 Rat2 Rat3 Rat4 Rat6 |
| P16636 | Protein-lysine 6-oxidase | Rat1 Rat2 Rat4 Rat5 Rat6 | Rat1 Rat2 Rat3 Rat4 Rat5 |
| D3Z9U8 | S100 calcium binding protein A7 like 2 | Rat1 Rat3 Rat4 Rat5 Rat6 | Rat1 Rat2 Rat3 Rat4 Rat5 Rat6 |
| P35053 | Glypican-1 | Rat2 Rat3 Rat4 Rat5 Rat6 | Rat1 Rat2 Rat3 Rat4 Rat5 Rat6 |
| Q642B0 | Glypican 4 | Rat2 Rat3 Rat4 Rat5 Rat6 | Rat1 Rat2 Rat3 Rat4 Rat5 Rat6 |
| P14668 | Annexin A5 | Rat1 Rat2 Rat3 Rat4 Rat5 | - |
| Q4G075 | Serine protease inhibitor EIA | Rat1 Rat2 Rat3 Rat4 Rat5 | - |
| Q9JJ19 | (Na+)/(H+)exchange regulatory cofactor NHE-RF1(NHERF-1) | Rat1 Rat2 Rat3 Rat4 Rat6 | - |
| A0A096MJ68 | Secretory leukocyte peptidase inhibitor | Rat1 Rat2 Rat3 Rat4 Rat6 | - |
| P17559 | Uteroglobin | Rat1 Rat2 Rat3 Rat5 Rat6 | - |
| P22282 | Cystatin-related protein 1 | Rat1 Rat2 Rat3 Rat5 Rat6 | - |
| P29975 | Aquaporin-1 | Rat1 Rat2 Rat4 Rat5 Rat6 | - |
| M0RCH6 | Charged multivesicular body protein 4B | Rat1 Rat2 Rat4 Rat5 Rat6 | - |
| D3ZLA3 | Copine 3 | Rat1 Rat2 Rat4 Rat5 Rat6 | - |
| Q6P6Q2 | Keratin, type II cytoskeletal 5 | Rat1 Rat3 Rat4 Rat5 Rat6 | - |
| P13265 | Glypican-3 | - | Rat1 Rat2 Rat3 Rat4 Rat5 Rat6 |
| Q30KJ2 | Beta-defensin 50 | - | Rat1 Rat2 Rat3 Rat4 Rat5 Rat6 |
| G3V7C6 | Tubulin beta chain | - | Rat1 Rat2 Rat3 Rat4 Rat5 Rat6 |
| D3ZAU0 | Mucin 5B, oligomeric mucus/gel-forming | - | Rat1 Rat2 Rat3 Rat4 Rat5 Rat6 |
| P51635 | Aldo-keto reductase family 1 member A1 | - | Rat1 Rat2 Rat3 Rat4 Rat5 Rat6 |
| P27590 | Uromodulin | - | Rat1 Rat2 Rat3 Rat4 Rat5 Rat6 |
| Q8VD89 | Ribonuclease pancreatic gamma-type | - | Rat1 Rat2 Rat3 Rat4 Rat5 Rat6 |
| A0A0G2JSZ3 | Neuroblastoma suppressor of tumorigenicity 1 | - | Rat1 Rat2 Rat3 Rat4 Rat5 Rat6 |
| P15399 | Probasin | - | Rat1 Rat2 Rat3 Rat4 Rat5 |
| F1LUS1 | Ig-like domain-containing protein | - | Rat1 Rat2 Rat3 Rat4 Rat5 |
| D4A0W2 | Lysozyme f1 | - | Rat1 Rat2 Rat3 Rat4 Rat5 |
| G3V8X4 | Gliomedin | - | Rat1 Rat2 Rat3 Rat4 Rat5 |
| Q5XI77 | Annexin | - | Rat1 Rat2 Rat3 Rat4 Rat6 |
| F1LRE2 | Insulin-like growth factor binding protein, acid labile subunit | - | Rat1 Rat2 Rat3 Rat4 Rat6 |
| A0A0G2K7Y9 | Zymogen granule protein 16B | - | Rat1 Rat2 Rat3 Rat4 Rat6 |
| P07861 | Neprilysin | - | Rat1 Rat2 Rat3 Rat4 Rat6 |
| A0A0G2QC50 | CD55 molecule | - | Rat1 Rat2 Rat3 Rat4 Rat6 |
| Q80VT8 | S100 calcium-binding protein | - | Rat1 Rat2 Rat3 Rat4 Rat6 |
| P02781 | Prostatic steroid-binding protein C2 | - | Rat1 Rat2 Rat3 Rat4 Rat6 |
| A0A0G2K2V6 | Keratin 10 | - | Rat1 Rat2 Rat3 Rat5 Rat6 |
| A0A0G2JWX4 | Keratin 2 | - | Rat1 Rat2 Rat3 Rat5 Rat6 |
| Q9QZH0 | Frizzled-4 | - | Rat1 Rat2 Rat3 Rat5 Rat6 |
| D4A5L9 | Similar to Cytochrome c | - | Rat1 Rat2 Rat3 Rat5 Rat6 |
| Q80YV8 | Stromal cell-derived factor 1 | - | Rat1 Rat2 Rat4 Rat5 Rat6 |
| Q6RUV5 | Ras-related C3 botulinum toxin substrate 1 | - | Rat1 Rat2 Rat4 Rat5 Rat6 |
| M0R5R0 | Protein S | - | Rat1 Rat2 Rat4 Rat5 Rat6 |
| B2RZB5 | Charged multivesicular body protein 2A | - | Rat1 Rat2 Rat4 Rat5 Rat6 |
| P61589 | Transforming protein RhoA | - | Rat1 Rat2 Rat4 Rat5 Rat6 |
| Q5M8C6 | Fibrinogen-like protein 1 | - | Rat1 Rat3 Rat4 Rat5 Rat6 |
| A0A0G2JV31 | X-prolyl aminopeptidase 1 | - | Rat1 Rat3 Rat4 Rat5 Rat6 |
| P00762 | Serine protease 1 | - | Rat1 Rat3 Rat4 Rat5 Rat6 |
| D4A281 | TNF receptor superfamily member 13C | - | Rat2 Rat3 Rat4 Rat5 Rat6 |
| P05964 | Protein S100-A6 | - | Rat2 Rat3 Rat4 Rat5 Rat6 |
| P02631 | Oncomodulin | - | Rat2 Rat3 Rat4 Rat5 Rat6 |
| A0A0G2JSG6 | Adenylate kinase 2 | - | Rat2 Rat3 Rat4 Rat5 Rat6 |
